# Voltage-Sensing Phosphatase (VSP) Regulates Endocytosis-Dependent Nutrient Absorption in Chordate Enterocytes

**DOI:** 10.1101/2022.03.10.483738

**Authors:** Adisorn Ratanayotha, Makoto Matsuda, Yukiko Kimura, Fumiko Takenaga, Tomoaki Mizuno, Md. Israil Hossain, Shin-ichi Higashijima, Takafumi Kawai, Michio Ogasawara, Yasushi Okamura

## Abstract

Voltage-sensing phosphatase (VSP) is a unique membrane protein that translates membrane electrical activities into the changes of phosphoinositide profiles. VSP orthologs from various species have been intensively investigated toward their biophysical properties, primarily using a heterologous expression system. In contrast, the physiological role of VSP in native tissues remains largely unknown. Here we report that zebrafish VSP (Dr-VSP) is functionally expressed on the endomembranes of lysosome-rich enterocytes (LREs) that mediate dietary protein absorption via endocytosis in the zebrafish mid-intestine. Dr-VSP-deficient LREs were remarkably defective in forming endosomal vacuoles after initial uptake of dextran and mCherry. Dr-VSP-deficient zebrafish exhibited growth restriction and higher mortality during the critical period when zebrafish larvae rely primarily on exogenous feeding via intestinal absorption. Furthermore, our comparative study on marine invertebrate *Ciona intestinalis* VSP (Ci-VSP) revealed co-expression with endocytosis-associated genes in absorptive epithelial cells of the *Ciona* digestive tract, corresponding to zebrafish LREs. These findings signify a crucial role of VSP in regulating endocytosis-dependent nutrient absorption in specialized enterocytes across animal species.

**Summary statement:** Voltage-sensing phosphatase (VSP) is identified in absorptive enterocytes, revealing its crucial role in promoting endocytosis and nutrient absorption during early development.

## Introduction

Phosphoinositides (PIPs) are essential phospholipids that constitute eukaryotic biological membranes. Differential phosphorylation of inositol head groups at the 3-, 4-, and 5-phosphate positions results in seven PIP species: three mono-phosphorylated PIPs, three bi-phosphorylated PIPs, and a single tri-phosphorylated PIP. These interconvertible PIP species are distinctly localized at the plasma membrane and subcellular compartments, regulating specific membrane-associated activities within the cells, including membrane dynamics, and cellular trafficking (Di Paolo and De Camilli, 2006; Falkenburger et al., 2010; Jones et al., 2013; Li et al., 2013; Wallroth and Haucke, 2018). Because of their essential roles, PIPs have attracted research attention for their regulatory mechanism and fundamental interplay in cell physiology.

Voltage-sensing phosphatase (VSP) is among the key molecules that regulate PIPs’ homeostasis. VSP is a unique membrane with two functional domains: the voltage sensor domain as typically found in voltage-gated ion channels; and the cytoplasmic catalytic region sharing molecular similarity to the phosphatase and tensin homolog deleted on chromosome 10 (PTEN), a tumor-suppressing PIP phosphatase. VSP functions upon membrane depolarization to exhibit voltage-dependent phosphatase activity towards PIPs; thus, directly translating membrane electrical activities into intracellular PIP signals. Unlike PTEN which exhibits rigid 3-phosphatase activity toward PI(3,4,5)P_3_ and PI(3,4)P_2_, VSP exhibits activities of both 3- and 5-phosphatases, thereby mediating four subreactions: dephosphorylating PI(3,4,5)P_3_ to PI(4,5)P_2_ or PI(3,4)P_2_; and from PI(4,5)P_2_ or PI(3,4)P_2_ to PI(4)P (Murata et al., 2005; Okamura et al., 2009; Okamura et al., 2018). Among four subreactions, the reaction with PI(4,5)P_2_ to produce PI(4)P is the most robust (Keum et al., 2016; Okamura et al., 2018). VSP orthologs are widely conserved across animal species, and their biophysical properties have been intensively investigated using in vitro approach. VSP expression has been detected in various animal tissues, including testis (Wu et al., 2001; Tapparel et al., 2003; Kawai et al., 2019), epithelium of gastrointestinal tract and renal tubules (Neuhaus and Hollemann, 2009; Ogasawara et al., 2011; Mutua et al., 2014; Ratzan et al., 2019), neurons (Murata et al., 2005; Yamaguchi et al., 2014a) and blood cells (Ogasawara et al., 2011). Our recent study has identified that mouse VSP is required for normal sperm motility through its role in regulating the spatial distribution of PI(4,5)P_2_ in sperm flagellum (Kawai et al., 2019). However, the physiological function of VSP in most organs remains elusive.

Previous studies demonstrated that zebrafish *Danio rerio* VSP (Dr-VSP) shares similar molecular architecture to other known VSP proteins and preserves their key biophysical properties (Hossain et al., 2008; Kawanabe et al., 2020). Zebrafish shows conserved physiology and anatomy with mammals (Gut et al., 2017). Its optical transparency during early development enables non-invasive monitoring and in vivo analysis of cellular behaviors (Kimmel et al., 1995; Vacaru et al., 2014). By taking these advantages of the zebrafish, we studied the physiological functions of VSP at the whole animal level. We show that VSP is highly expressed in lysosome-rich enterocytes (LREs), which are specialized enterocytes that contain large supranuclear vacuoles and absorb dietary protein from the intestinal lumen (Iwai, 1969; Stroband and van der Veen, 1981; Rombout et al., 1985; Park et al., 2019). We present evidence from in vivo assays that VSP contributes to the endocytosis of enterocytes. Comparative analysis on ascidian *Ciona intestinalis* Type A also reveals co-expression of VSP (Ci-VSP) with the endocytic adaptor Dab2 (Ci-Dab2) in absorptive epithelial cells of the digestive tract, suggesting that the function of VSP in enterocytes is evolutionarily conserved. These results highlight a crucial role of VSP in the endocytosis-dependent nutrient absorption of intestinal epithelial cells.

## Results

### 1. Dr-VSP is expressed in the zebrafish intestine

RT-PCR analysis revealed substantial expression of Dr-VSP mRNA in testis, spleen, intestine, kidney, and gills of adult zebrafish (Figure 1A). Dr-VSP mRNA was also detected in 7-dpf zebrafish larvae (Figure 1B). Whole-mounted in situ hybridization (WISH) showed specific expression of Dr-VSP mRNA in the mid-intestine (Figure 1C), with the signal detectable as early as 5 days after fertilization (dpf). The result was comparable to the EGFP signal observed in our *Tg(vsp:EGFP)* transgenic zebrafish (Figure 1, D and E), which were generated using CRISPR-Cas9-mediated knock-in transgenesis (Kimura et al., 2014) to express EGFP recapitulating endogenous expression of *vsp*, the Dr-VSP-encoding gene (Supplementary Figure S1). These findings suggest a potential role of Dr-VSP in the digestive tract from an early developmental stage.

**Figure 1:**
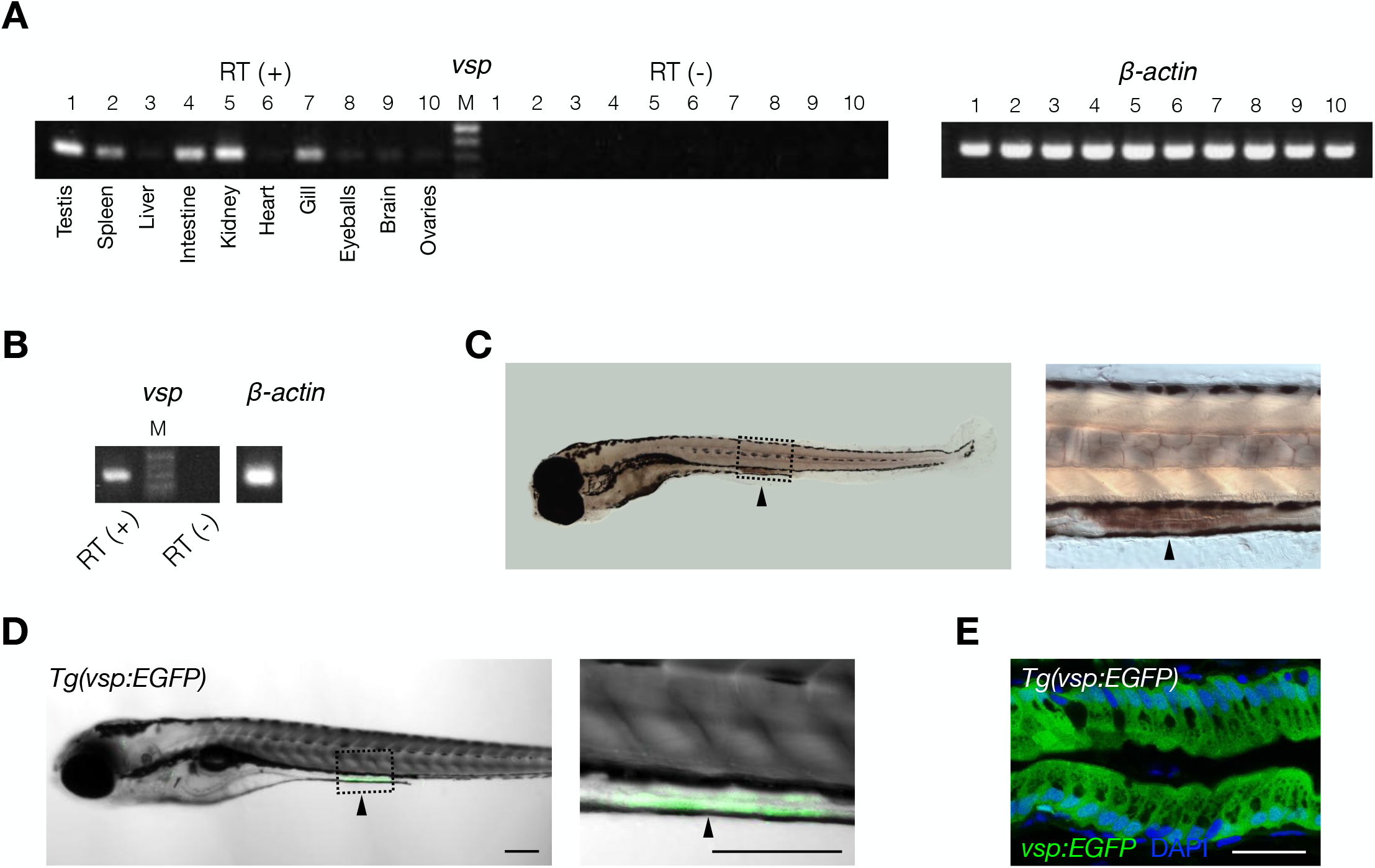
Expression profile of the Dr-VSP-encoding gene (*vsp*) in zebrafish tissues. (A) RT (+) *vsp* expression in adult zebrafish tissues: (1) testis, (2) spleen, (3) liver, (4) intestine, (5) kidney, (6) heart, (7) gill, (8) eyeballs, (9) brain, (10) ovaries. RT (-), negative control for all tissues in the same order. β*-actin*, positive control. M, marker. (B) RT (+), *vsp* expression in 7-dpf zebrafish larva. RT (-), negative control. β*-actin*, positive control. M, marker. (C) Whole-mounted in situ hybridization (WISH) of 6-dpf zebrafish larva showing *vsp*-positive signal (arrowhead) in enterocytes of mid-intestine. (D) A 5-dpf *Tg(vsp:EGFP)* zebrafish larva expressing EGFP under the promotor of the Dr-VSP-encoding gene in enterocytes of mid-intestine (arrowhead). The EGFP-positive cells in this figure correspond to the *vsp*-positive cells in Figure 1C. Green, Dr-VSP (*vsp:EGFP*). Scale bar = 200 µm. (E) Confocal image of larval enterocytes at mid-intestine of a 5-dpf *Tg(vsp:EGFP)* zebrafish larva (sagittal section). The enterocytes in this region are known as lysosome-rich enterocytes (LREs) that contain large supranuclear vacuoles. Green, Dr-VSP (*vsp:EGFP*). Blue, DAPI. Scale bar = 25 µm

### 2. Dr-VSP is spatially distributed inside zebrafish enterocytes

Dr-VSP protein expression was examined using an anti-Dr-VSP antibody (NeuroMab clone N432/21). The immunostainings detected abundant expression of Dr-VSP protein in the epithelial cells of larval pronephros and posterior portion of the mid-intestine (Figure 2A), correlating with the results of WISH and *Tg(vsp:EGFP)* transgenesis.

**Figure 2:**
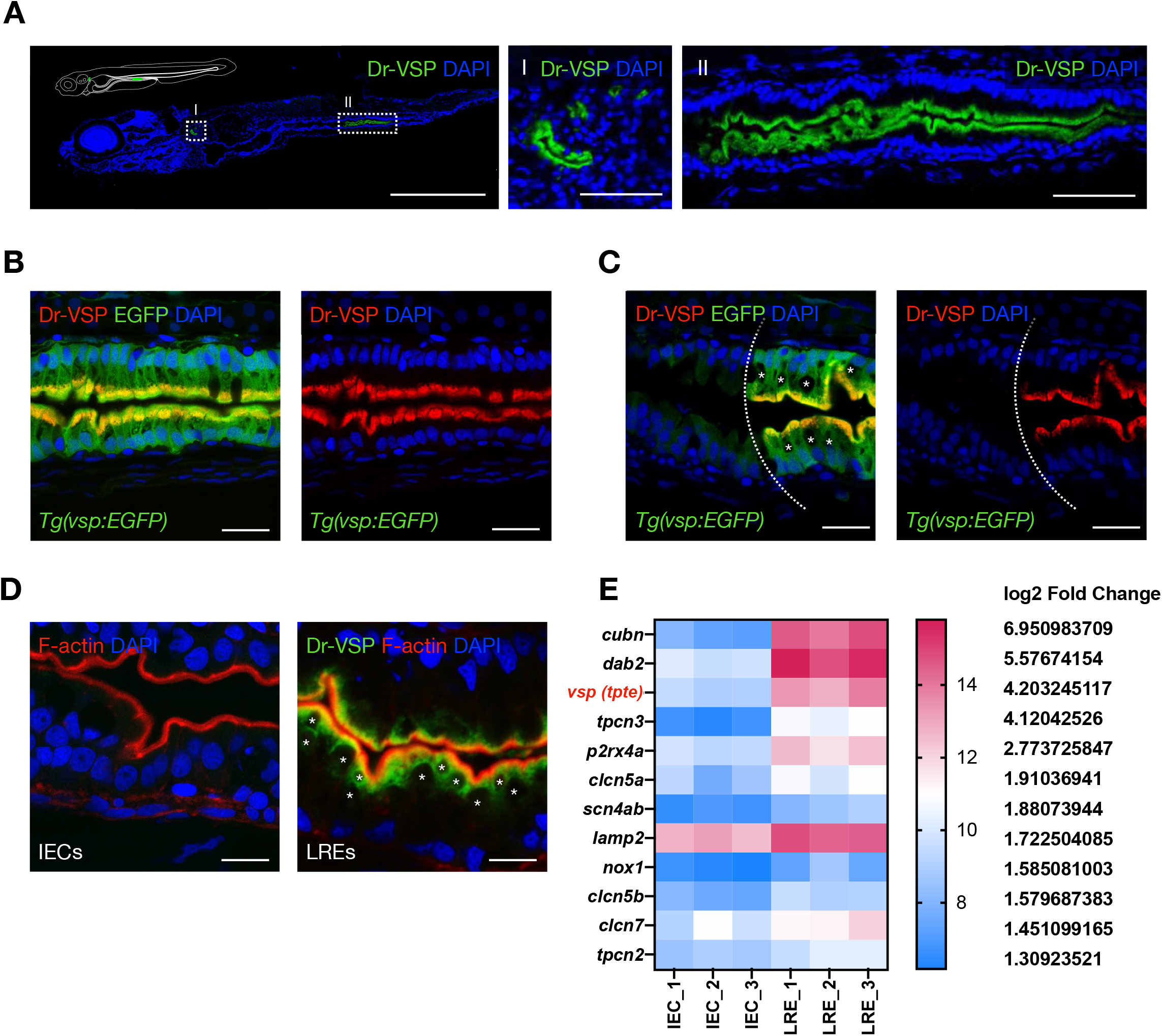
Expression profile Dr-VSP protein in zebrafish tissues. (A) Confocal images of Dr-VSP immunostaining in a 7-dpf wild-type zebrafish larva (sagittal section). Dr-VSP is highly expressed in (I) larval pronephros and (II) enterocytes at the posterior part of mid-intestine, which are defined as lysosome-rich enterocytes (LREs). Green, Dr-VSP. Blue, DAPI. Scale bar = 500 µm (left) and 50 µm (middle and right) (B) Dr-VSP immunostaining in LREs of a 7-dpf *Tg(vsp:EGFP)* transgenic zebrafish larva (sagittal section). Dr-VSP expression signals were completely co-localized with EGFP signals in LREs. Green, Dr-VSP *(vsp:EGFP)*. Red, Dr-VSP (antibody). Blue, DAPI. Scale bar = 20 µm. (C) Dr-VSP immunostaining in a 7-dpf *Tg(vsp:EGFP)* transgenic zebrafish larva (sagittal section), showing a border between intestinal epithelial cells (IECs; VSP-negative) and LREs (VSP-positive). LREs contain large supranuclear vacuoles in the cytoplasm, contrasting to IECs. Green, Dr-VSP *(vsp:EGFP)*. Red, Dr-VSP (antibody). Blue, DAPI. Scale bar = 20 µm. Background signal of EGFP channel was increased in the left figure to enhance visibility of IECs. (D) Dr-VSP immunostaining in IECs (left) and LREs (right) of a 7-dpf wild-type zebrafish larva (sagittal section). Dr-VSP expression was restricted mainly to LREs, but not detectable in other IECs. Green, Dr-VSP. Red, F-actin. Blue, DAPI. Scale bar = 10 µm. Images were visualized under LSM880 confocal microscope with Airy Scan. (E) Heatmap of RNA-seq data representing expression levels of selected genes in IECs and LREs. Raw data of RNA-seq experiments were retrieved from the GEO Datasets database with accession number GSE124970 (Park et al., 2019), and analyzed using web-based application iDEP v0.92 (Ge et al., 2018).

Here we focused on Dr-VSP expression in the intestine. Principal enterocytes in this region have been characterized as lysosome-rich enterocytes (LREs) that mediate dietary protein absorption via endocytosis (Iwai, 1969; Stroband and van der Veen, 1981; Rombout et al., 1985; Park et al., 2019). Dr-VSP immunostaining in the intestine of *Tg(vsp:EGFP)* transgenic zebrafish revealed complete co-localization of Dr-VSP signal and EGFP-positive LREs (Figure 2B). Notably, Dr-VSP expression is absent in the intestinal epithelial cells (IECs), which morphologically differ from LREs by the absence of large supranuclear vacuoles (Figure 2, C and D) (Ng et al., 2005). Transcriptome database of zebrafish LREs (Figure 2E) also supports that Dr-VSP is highly enriched in LREs, along with other genes known to be associated with macromolecule absorption, such as *lamp2, cubn*, and *dab2* (Rodriguez-Fraticelli et al., 2015; Park et al., 2019).

The positive signal of the anti-Dr-VSP antibody was not entirely co-localized with phalloidin staining, which represents actin filaments (F-actin) in microvilli (Figure 3, A and B), indicating that Dr-VSP in LREs is preferentially distributed at intracellular region beneath the apical surface rather than at the cell surface or microvilli. Quantification of fluorescence (Figure 3C) demonstrates that Dr-VSP showed the most intense signal at a deeper region of the cells than the microvillous F-actins. Immuno-electron microscopy analysis revealed a particularly high density of immunoparticles at the subapical region of LREs, while it is rarely detectable at microvilli (Figure 4A, Supplementary Figure S2), corresponding to the immunofluorescence results. To specify which intracellular structures contain Dr-VSP, we used the MDCKII cells as a model for examining the intracellular compartments involved in epithelial endocytosis (Parton, 1992), simulating zebrafish LRE function. In MDCKII cells, Dr-VSP was co-expressed with (i) Rab5, an early endosome marker; or (ii) Rab11, a recycling endosome marker. The Dr-VSP signal was broadly co-localized with Rab5-positive and Rab11-positive intracellular compartments (Figure 4B, Supplementary Movie S1, Supplementary Movie S2), indicating that Dr-VSP is expressed on the membranes of early and recycling endosomes.

**Figure 3:**
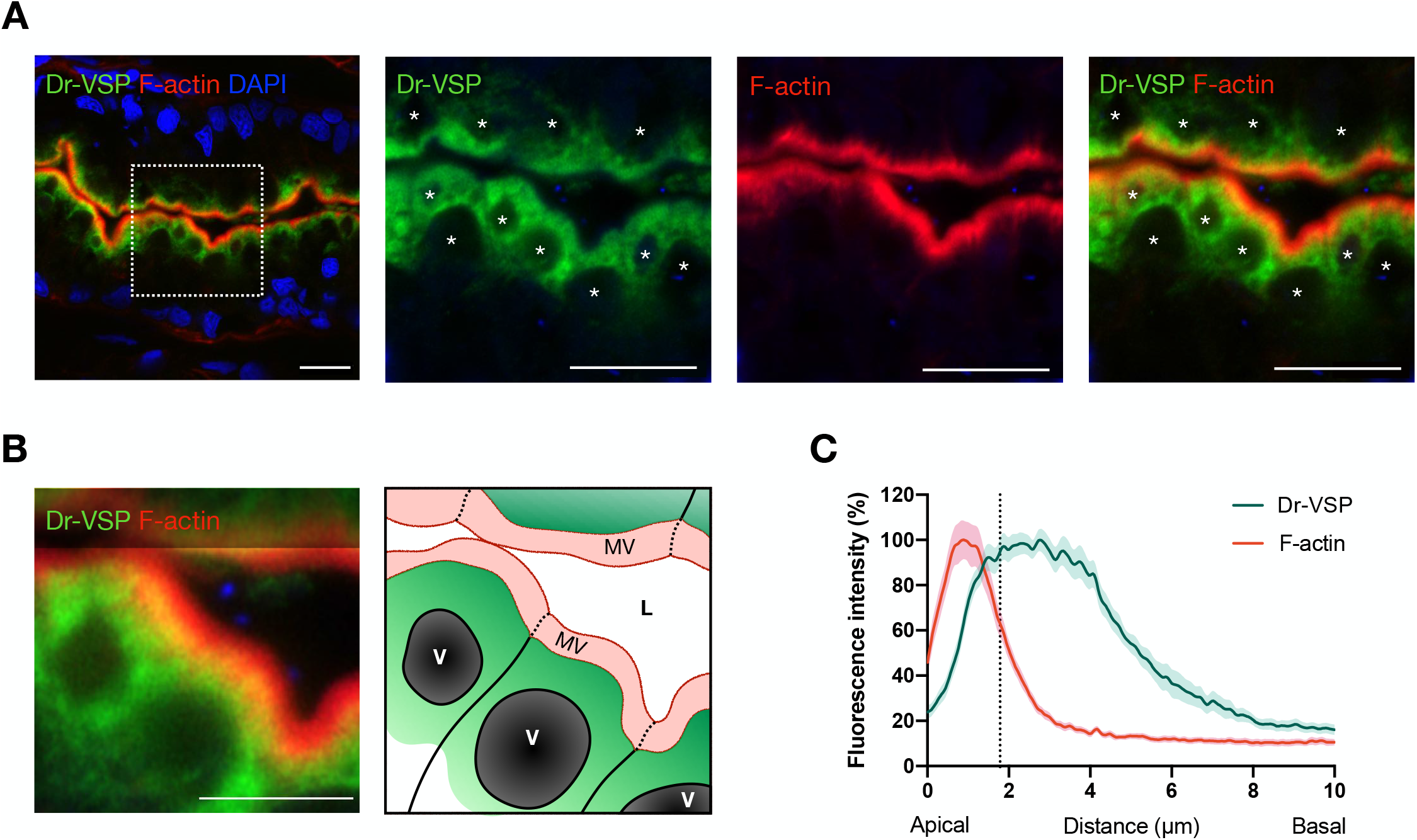
Expression profile Dr-VSP protein at subapical region of LREs. (A) Confocal images of Dr-VSP immunostaining showing the subcellular distribution of Dr-VSP in larval LREs. Dr-VSP is primarily localized in the cytoplasm underneath apical surface, but barely detectable at the microvilli and plasma membrane of LREs. Green, Dr-VSP. Red, F-actin, Blue, DAPI. *Vacuole. Scale bar = 10 µm. (B) Confocal image of larval LREs in a higher magnification and their corresponding schematic illustrations, focusing on the intracellular distribution of Dr-VSP. Green, Dr-VSP. Red, F-actin. Blue, DAPI. V, vacuole. MV, microvilli. L, lumen. Scale bar = 5 µm. (C) Fluorescence intensity of Dr-VSP (green) and F-actin (red) along the apico-basal axis of individual LREs. Data are presented as mean ± s.e.m. Enterocytes, ≥ 16 cells from 4 larvae for each zebrafish line. Dotted line in the graph conceptually indicates the junction between microvilli and cell body.

**Figure 4:**
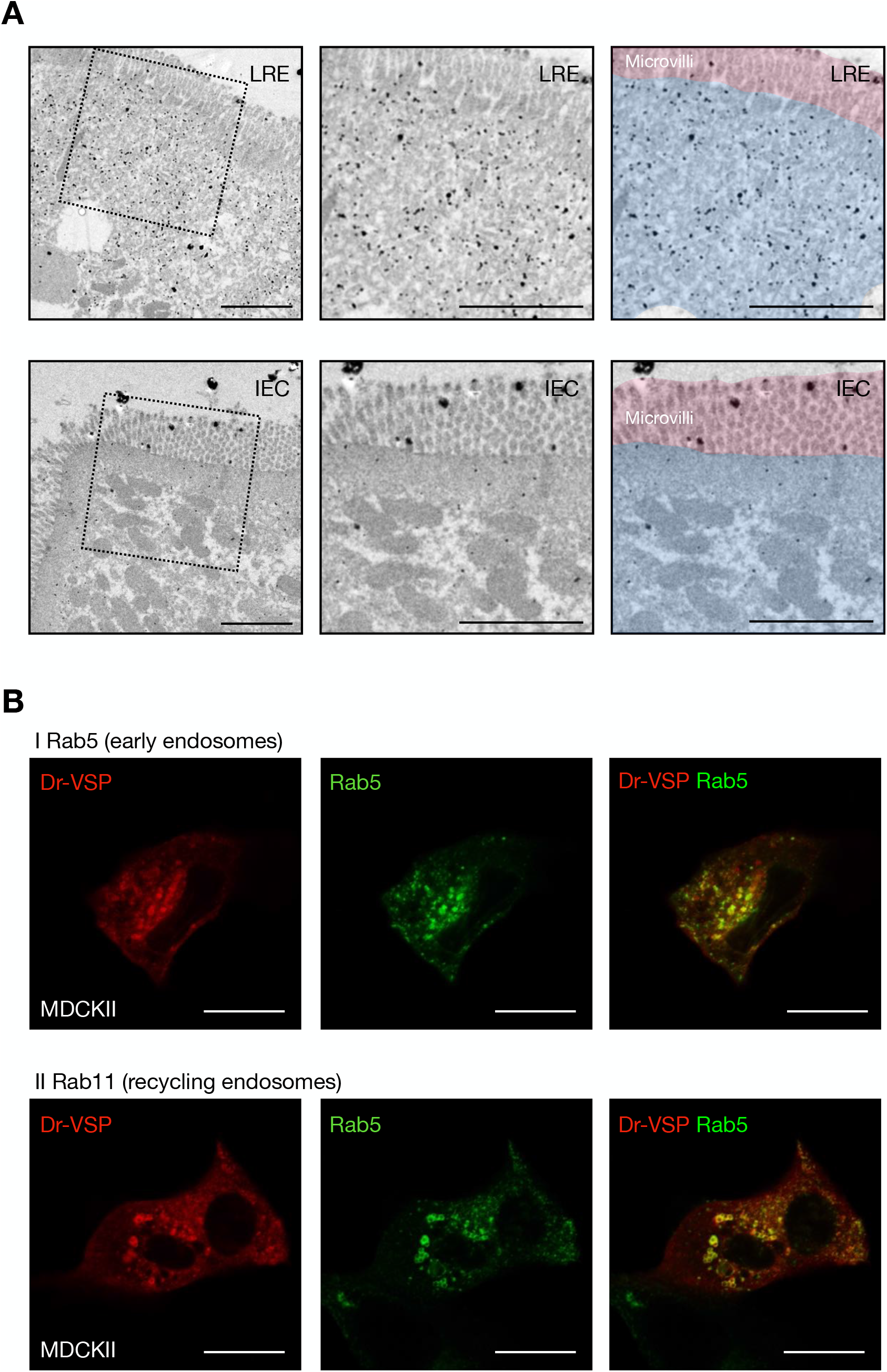
Expression profile Dr-VSP protein at subapical region of LREs. (A) Pre-embedding silver-enhanced immunogold staining for Dr-VSP in LREs (top) and IECs (bottom) of 14-dpf wild-type zebrafish larva. Right two images are the same as the middle images but were highlighted with red and blue for microvilli and subapical region, respectively. In LREs, immunoparticles for Dr-VSP are preferentially distributed at the subapical region while rarely detectable at microvilli, consistent with results of Dr-VSP immunofluorescence in Figure 3. In IECs, immunoparticles for Dr-VSP are rarely detectable at both subapical region and microvilli, consistent with Dr-VSP immunofluorescence and RNA-seq analysis in Figure 2. Scale bar = 2 µm. (B) MDCKII cells co-expressing Dr-VSP-mCherry and (I) Rab5-EGFP for early endosomes; or (II) Rab11-EGFP for recycling endosomes. Dr-VSP is localized at the endosomal membranes of early and recycling endosomes. Red, Dr-VSP. Green, Rab5 or Rab11. Scale bar = 20 µm.

### 3. Dr-VSP contributes to the survival and growth of zebrafish larvae

To investigate Dr-VSP function under physiological conditions, we generated Dr-VSP-deficient (Dr-VSP^-/-^) zebrafish using CRISPR-Cas9-mediated mutagenesis, targeting the early exon of the *vsp* gene. Mutant zebrafish lacking the transmembrane segments and cytoplasmic catalytic region of Dr-VSP were obtained (Supplementary Figure S3).

Dr-VSP^-/-^ larvae showed normal gross morphology and behavior. However, we noticed that the survival rate of Dr-VSP^-/-^ mutants was significantly lower than that of wild-type larvae at the age of 10 – 14 dpf (Figure 5A). During this period, the yolk is completely reabsorbed, and zebrafish growth is generally dependent on exogenous feeding via the digestive system (Ng et al., 2005; Wallace et al., 2005). Moreover, Dr-VSP^-/-^ mutants that survived from the early period exhibited long-term growth restriction, as evidenced by their smaller size compared to their sibling wild-types (Figure 5, B and C). Despite the absence of obvious congenital deformities, these results could be inferred that Dr-VSP function contributes to the survival and growth of zebrafish larvae during early life. The remaining Dr-VSP^-/-^ mutants thrived to adulthood and reached productive maturity at around the same age as wild-type zebrafish without apparent fertility defect.

**Figure 5:**
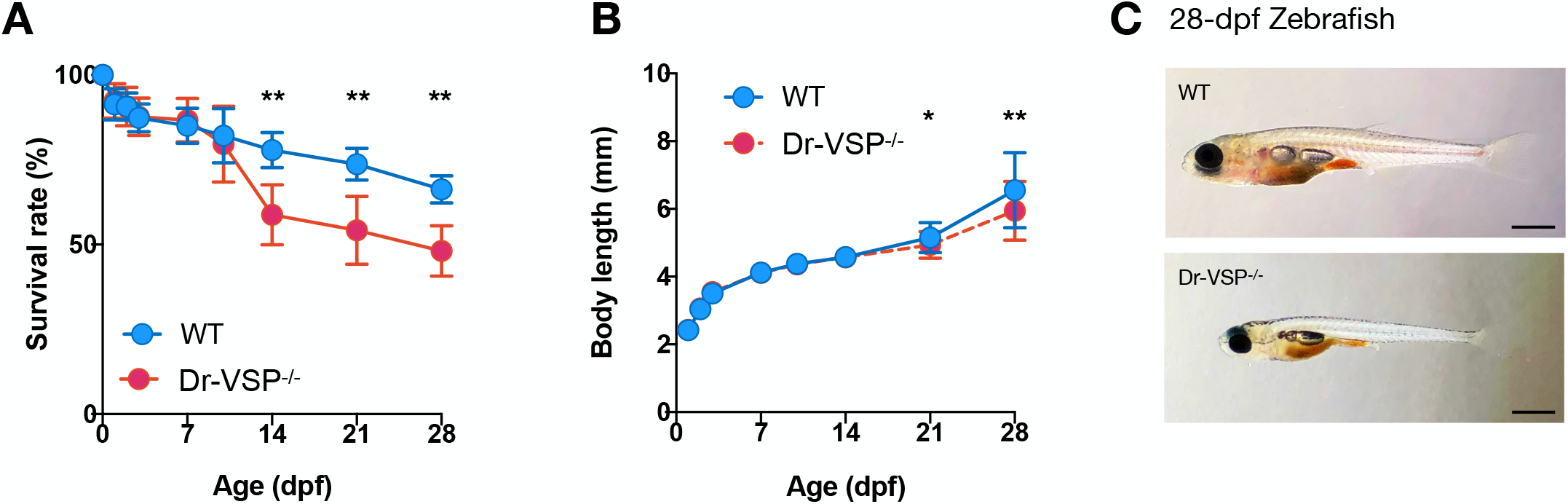
Phenotypic investigation of Dr-VSP^-/-^ zebrafish. (A) Survival rates of wild-type and Dr-VSP^-/-^ zebrafish larvae during the first month of age. Zebrafish larvae, 300 at D0 for each zebrafish line. All statistical data were collected from three independent experiments. (B) Body length of wild-type and Dr-VSP^-/-^ zebrafish larvae raised under non-calorie restricted standard diet. Zebrafish larvae, ≥ 40 on the first day (D0) for each zebrafish line. Error bars, means ± s.d.; *P < 0.05; **P < 0.01; unpaired Student’s t-test. (C) Brightfield images of the representative wild-type (top) and Dr-VSP^-/-^ (bottom) zebrafish larvae at 28 dpf, raised under the same conditions. Scale bar = 1 mm.

### 4. Dr-VSP facilitates endocytic nutrient absorption in larval LREs

Dr-VSP is abundant at the endomembranes inside LREs, which absorb nutrients via endocytosis (Iwai, 1969; Stroband and van der Veen, 1981; Rombout et al., 1985; Park et al., 2019), suggesting that Dr-VSP could be involved in this specialized process. To test this hypothesis, we compared the endocytosis efficiency between the wild type and Dr-VSP^-/-^ larvae by gavaging 6-dpf zebrafish larvae with either fluorescent dextran (fDex) or mCherry solution directly into the digestive tract (Figure 6A) (Cocchiaro and Rawls, 2013), and subsequently observed endocytosis in LREs using confocal microscopy. fDex was used for assessing fluid-phase endocytosis; and mCherry was used as a protein cargo for assessing receptor-mediated endocytosis (Rodriguez-Fraticelli et al., 2015; Park et al., 2019).

**Figure 6:**
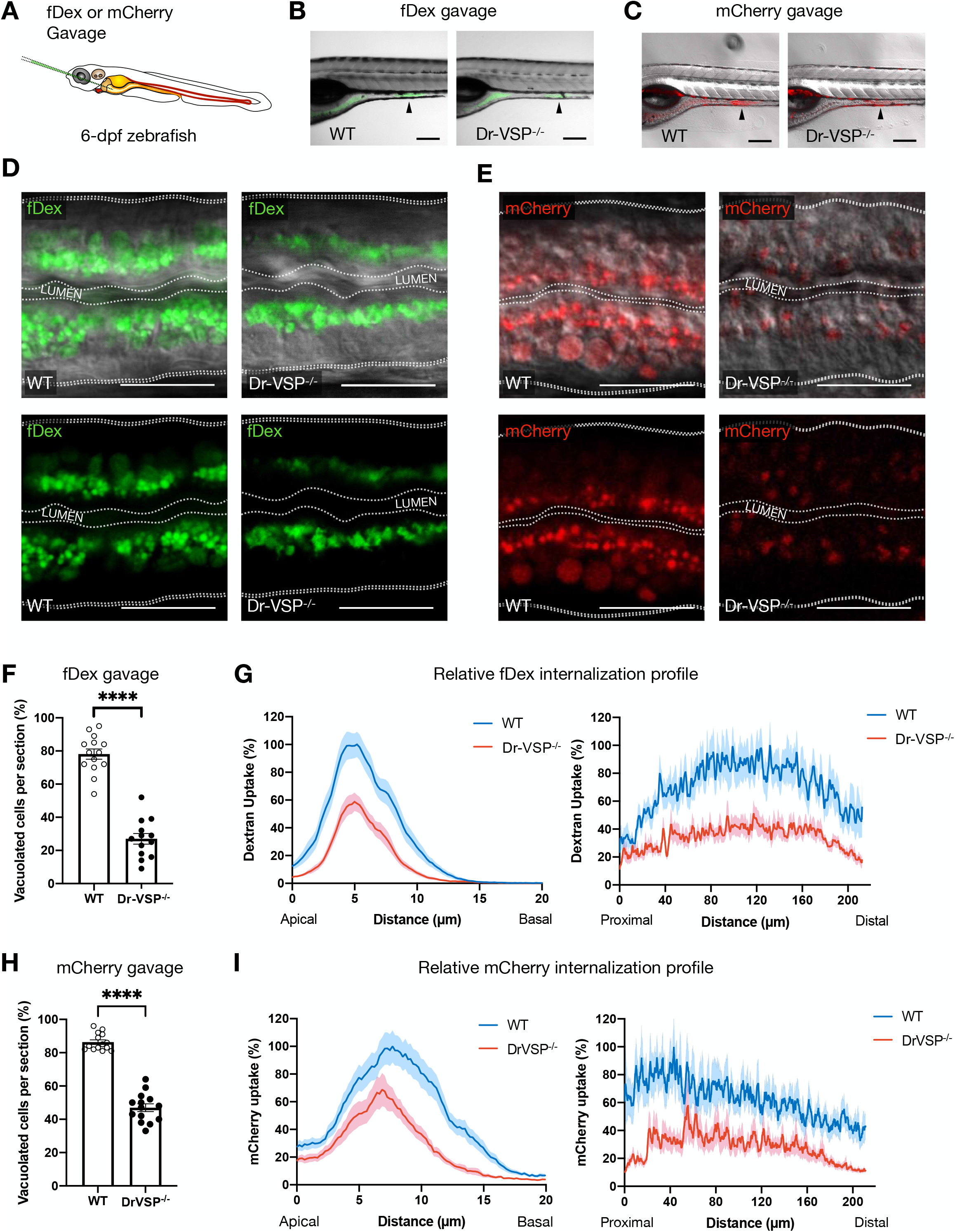
Functional analysis of Dr-VSP in larval enterocytes. (A) Experimental approach: 6-dpf zebrafish larvae were gavaged with Alexa Fluor™ 488-tagged dextran (fDex) or mCherry solutions into the anterior intestinal bulb. Live imaging was performed at 2 hours after gavage. (B – E) fDex and mCherry internationalized into zebrafish LREs after gavage. (B) Live confocal images of wild-type (WT; left) and Dr-VSP^-/-^ (right) zebrafish larvae showing fDex internalization in enterocytes at the posterior portion of mid-intestine (arrowhead). Scale bar = 200 µm (C) Live confocal images of wild-type (WT; left) and Dr-VSP^-/-^ (right) zebrafish larvae showing mCherry internalization in enterocytes at the posterior portion of mid-intestine (arrowhead). Scale bar = 200 µm (D) Live confocal images of wild-type (left) and Dr-VSP^-/-^ (right) enterocytes showing fDex internalization. The images were presented with DIC (top) or without DIC (bottom) to enhance visibility of the internalized fDex. Dotted lines conceptually indicate the outlines of larval intestines. Scale bar = 20 µm (E) Live confocal images of wild-type (left) and Dr-VSP^-/-^ (right) enterocytes showing mCherry internalization. The images were presented with DIC (top) or without DIC (bottom) to enhance visibility of the internalized mCherry. Dotted lines conceptually indicate the outlines of larval intestines. Scale bar = 20 µm (F and G) Endocytosis efficiency analysis in LREs after fDex gavage. (F) Percentage of the enterocytes containing fDex-filled lysosomal vacuoles at 2 hours following gavage. Error bars, means ± s.e.m.; ****P < 0.0001; unpaired Student’s t-test. Number of enterocytes = 344 cells from 14 wild-type and 359 cells from 13 Dr-VSP^-/-^ zebrafish. (G) Relative fDex internalization profile comparing between wild-type and Dr-VSP^-/-^ zebrafish larvae at 2 hours following gavage. Data are means ± s.e.m. percentage of intracellular fDex intensity along the specified axes, with a 100% indicates the maximal value of the average intensity on each axis. (Left) Apico-basal axis of individual cells. (Right) Longitudinal axis along the posterior portion of mid-intestine. Number of fish = 14 for wild-type and 13 for Dr-VSP^-/-^. Data are presented as mean ± s.e.m., collected from 4 independent experiments for both wild-type and Dr-VSP^-/-^. (H and I) Endocytosis efficiency analysis in LREs after mCherry gavage. (H) Percentage of the enterocytes containing mCherry-filled lysosomal vacuoles at 2 hours following gavage. Error bars, means ± s.e.m.; ****P < 0.0001; unpaired Student’s t-test. Number of enterocytes = 376 cells from 14 wild-type and 341 cells from 14 Dr-VSP^-/-^ zebrafish. (I) Relative mCherry internalization profile comparing between wild-type and Dr-VSP^-/-^ zebrafish larvae at 2 hours following gavage. Data are means ± s.e.m. percentage of intracellular mCherry intensity along the specified axes, with a 100% indicates the maximal value of the average intensity on each axis. (Left) Apico-basal axis of individual cells. (Right) Longitudinal axis along the posterior portion of mid-intestine. Number of fish = 14 for each zebrafish line. Data are presented as mean ± s.e.m., collected from 4 independent experiments for both wild-type and Dr-VSP^-/-^.

At two hours after gavage, fDex and mCherry were absorbed into LREs in both wild-type and Dr-VSP^-/-^ zebrafish larvae (Figure 6, B and C). However, the uptake of fDex and mCherry were reduced in Dr-VSP^-/-^ LREs (Figure 6, D and E). In the experiment with fDex, the fluorescent signals in Dr-VSP^-/-^ LREs were mostly confined to apical vesicles, with less signal found in larger endosomes and supranuclear vacuoles (Figure 6D). The average percentage of LREs containing fDex-filled endosomes was nearly 80% in wild-types but less than 30% in Dr-VSP^-/-^ larvae (Figure 6F). We further analyzed the relative fDex internalization profile of LREs by measuring fluorescent intensity within the cells (Figure 6G) (Rodriguez-Fraticelli et al., 2015; Park et al., 2019). As shown in Figure 6G, fDex uptake into Dr-VSP^-/-^ LREs was clearly impaired both in the apicobasal axis of individual cells and the anteroposterior axis across longitudinal segments of LREs. Similar results were obtained in the mCherry experiment: the average percentage of LREs containing mCherry-filled endosomes was greater than 80% in wild-types versus 40% in Dr-VSP^-/-^ larvae (Figure 6H). Spatial profile of the marker distribution in the cells (Figure 6I) also showed reduced uptake of mCherry in Dr-VSP^-/-^ LREs.

When we gavaged wild-type zebrafish with the mixture of fDex and mCherry, we found that while most LREs contained both fluorescent cargoes in the same supranuclear vacuoles, some vacuoles contained only fDex or mCherry (Supplementary Figure S4A). In Dr-VSP^-/-^ mutants, on the other hand, mCherry was transported into some supranuclear vacuoles, whereas fDex was mostly found in apical vesicles inside Dr-VSP^-/-^ LREs (Supplementary Figure S4B). This suggests that the uptake defect in Dr-VSP^-/-^ LREs was more severe with fDex than with mCherry.

Taken together, these findings support our hypothesis that Dr-VSP is involved in the endocytosis of LREs, specifically in the maturation of endosomal vacuoles following initial endocytosis. The findings also suggest that Dr-VSP plays a greater role in fluid-phase endocytosis, the primary mechanism of fDex uptake into cells.

### 5. Dr-VSP maintains intact LRE morphology

We examined the LRE morphology of 14-dpf zebrafish larvae, the age around one week after zebrafish enterocytes completed morphogenesis (Figure 7A) (Ng et al., 2005; Wallace et al., 2005). This age also corresponds to the period when higher mortality was observed in Dr-VSP^-/-^ larvae (Figure 5A). We measured key parameters that represent cell morphology: central height, apical width, basal width, and microvillus length (Sidhaye et al., 2016). The measurement revealed that Dr-VSP^-/-^ LREs were significantly shorter in all other parameters except for the microvillus length (Figure 7B). The fact that Dr-VSP deficiency primarily affected the cell body is consistent with the idea that Dr-VSP plays a major role at the endomembranes within cells than at the plasma membrane of microvilli. The defect in LRE morphology was not observed at early stages, such as at 6 dpf (Figure 7C). We examined histological sections of 14-dpf zebrafish LREs with toluidine blue staining and analyzed endosomal vacuoles, comparing wild-type and Dr-VSP^-/-^ mutants (Figure 7D). Although supranuclear vacuoles could be seen in most cells, the measurement of vacuolar perimeter and area showed that their size was apparently smaller in Dr-VSP^-/-^ LREs (Figure 7, E and F). We also used transmission electron microscopy (TEM) to define the ultrastructures of 14-dpf LREs, focusing on the subapical region where Dr-VSP is typically expressed. TEM images of wild-type and Dr-VSP^-/-^ LREs (Figure 7G) revealed key features of absorptive enterocytes, such as membrane invaginations at inter-microvillous spaces, cytoplasmic tubules, branching tubules, and endocytic vesicles associated with invaginated membranes (tubule-vesicle complexes) (Iwai, 1969; Iida and Yamamoto, 1985; Iida et al., 1986).

**Figure 7:**
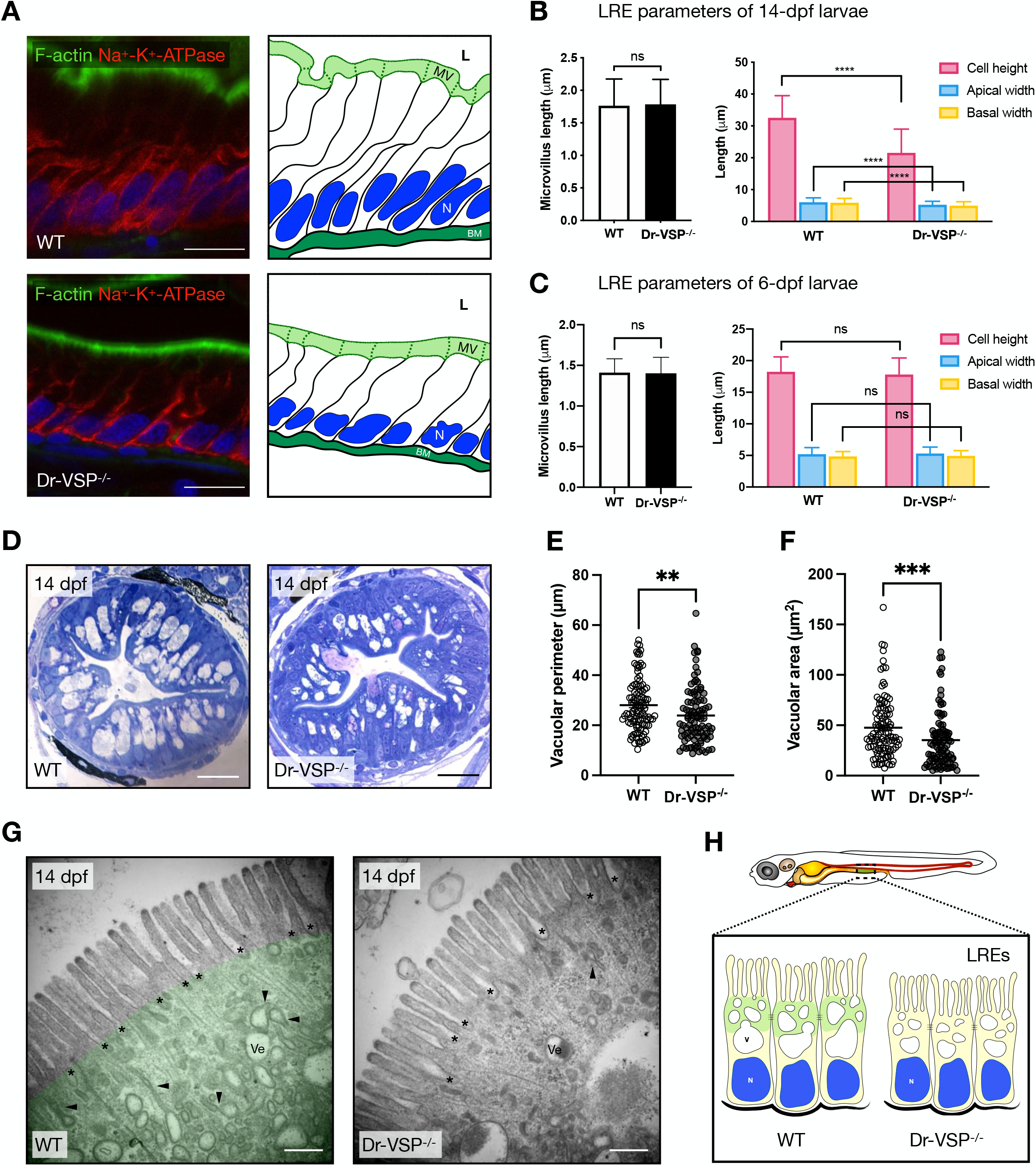
Morphological change in Dr-VSP^-/-^ LREs. (A) Immunostaining showing cellular framework in sagittal sections of 14-dpf wild-type (top) and Dr-VSP^-/-^ (bottom) LREs (left); and their corresponding schematic illustration (right). Green, F-actin. Red, Na^+^-K^+^ ATPase. Blue, DAPI. Scale bar = 10 µm. (B) Cellular parameters of LREs at 14 dpf comparing between wild-type and Dr-VSP^-/-^ zebrafish larvae. (Left) Microvillus length. (Right) Cell height, apical width, and basal width. Enterocytes, ≥ 120 cells from 5 larvae for each zebrafish line. (C) Cellular parameters of LREs at 6 dpf comparing between wild-type and Dr-VSP^-/-^ zebrafish larvae. (Left) Microvillus length. (Right) Cell height, apical width, and basal width. Enterocytes, ≥ 170 cells from 6 larvae for each zebrafish line. Error bars, means ± s.d.; ****P < 0.0001; unpaired Student’s t-test; ns, no statistically significant difference. (D) Representative transverse sections at mid-intestine of 14-dpf wild-type (left) and Dr-VSP^-/-^ (right) zebrafish larvae, showing LREs with intracellular supranulcear vacuoles. Sections were stained with toluidine blue. Scale bar = 20 µm. (E – F) Vacuole parameters of 14-dpf LREs. (E) Vacuolar perimeter of wild-type (WT) and Dr-VSP^-/-^ LREs. **P = 0.0024; Mann-Whitney *U* test. Vacuoles, 108 for wild-type and 95 for Dr-VSP^-/-^. Data were collected from 3 larvae for each zebrafish line. (F) Vacuolar area of wild-type (WT) and Dr-VSP^-/-^ LREs from the same samples as in Figure 6E. ***P = 0.0003; Mann-Whitney *U* test. (G) Representative TEM images at mid-intestine of 14-dpf wild-type (left) and Dr-VSP^-/-^ (right) zebrafish LREs, showing the ultrastructures of microvilli and subapical region. Key features of absorptive enterocytes are presented, including membrane invaginations at inter-microvillous spaces (*), cytoplasmic tubules and tubule-vacuole complexes (arrowhead), and numerous endocytic vesicles (Ve). In wild-type LRE, green represents the region upper to supranuclear vacuole, which corresponds to the area with positive immunofluorescence signal of Dr-VSP in Figure 3, A and B. Scale bar = 500 nm. (H) Schematic diagram of wild-type (WT) and Dr-VSP^-/-^ LREs. Dr-VSP (green) is primarily localized in cytoplasm close to the apical surface of enterocytes, promoting endosomal maturation during endocytosis-dependent nutrient absorption. N, nucleus. V, Vacuoles.

Interestingly, these features appeared less prominent in Dr-VSP^-/-^ LREs. Further observations suggested that small endocytic vesicles in Dr-VSP^-/-^ LREs are located at deeper regions than those in wild-types, increasing the distance between the apical surface and vesicle-dense area (Supplementary Figure S5). Moreover, branching tubules and tubule-vacuole complexes were less frequently present in the subapical region of mutant LREs (Supplementary Figure S6), seemingly reflecting the disturbance in endosome maturation. Collectively, these results suggest that LREs require Dr-VSP in regulating the formation and expansion of endosomal vacuoles, which in turn maintain intact LRE morphology. Figure 7H summarizes how Dr-VSP contributes to endosomal vacuoles and the morphology of LREs.

### 6. Endocytic role of VSP is potentially conserved in the ascidian digestive tract

VSP gene is widely conserved among chordates (Supplementary Figure S7, Supplementary Figure S8). To determine if the role of VSP in the endocytosis of enterocytes is conserved among animal species, we investigated the expression profile of sea squirt *C. intestinalis* VSP (Ci-VSP) in the digestive tract. Sea squirt belongs to the basal chordates, and the gene expression profiles of *C. intestinalis* Type A were studied in great detail with reference to chordate evolution (Nakayama et al., 2019). WISH on transparent 2-week-old *Ciona* juveniles revealed the expression of Ci-VSP mRNA in the stomach, mid-intestine, and posterior intestine (Figure 8A), which were known as absorptive regions of the *Ciona* digestive tract (Yonge, 1925). Higher magnification showed a strip-like expression pattern both in the stomach and bulged intestine. Similar patterns were found in the digestive tract of *Ciona* young adults, with the expression signal being more robust in the stomach and mid-intestine than in the posterior intestine (Figure 8B). Additional Ci-VSP expression was observed in the looped region of the intestine (Figure 8B, white arrowhead), which corresponds to a region expressing several absorptive genes; including peptide transporter (PEPT1) and monosaccharide transporters (SGLT1, GLUT5) (Supplementary Figure S9).

**Figure 8:**
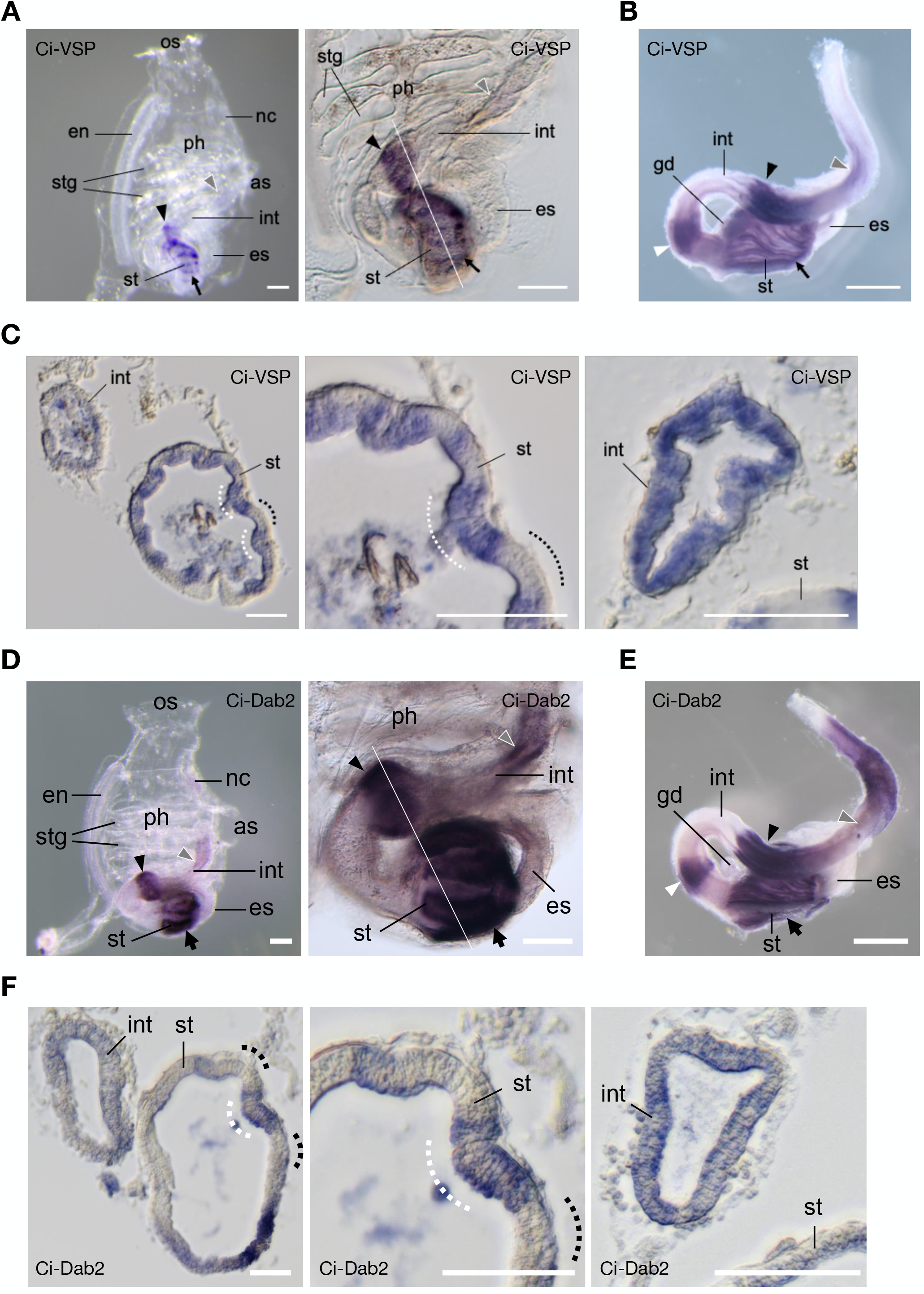
Expression profile of Ci-VSP and Ci-Dab2 in ascidian *C. intestinalis*. (A – C) Expression profile of Ci-VSP by WISH. (A) Whole-mount (left) 2-week-old juvenile showing regional expressions in the stomach (st; arrow), bulged-region of the mid-intestine (int; arrowhead), and posterior-intestine (gray arrowhead). Higher magnification (right) reveals a stripe-like distribution of signals in the stomach and bulged intestine. (B) Dissected post-pharyngeal region of the digestive tract of the 1.5-month-old young adult showing an additional expression region (white arrowhead) in the looped-intestine. (C) Transverse section specimens of the juvenile (white line in A) showing Ci-VSP expressions in the epithelium of the digestive tract (left). Expression signals in the stomach epithelium are restricted mainly in the inner-folds (white dotted lines), which contain several absorptive enterocytes (Hayashibe et al., 2017), as opposed to the outer-folds (dotted lines) with many pancreatic-related exocrine genes expressions (Nakayama and Ogasawara, 2017). Circumferences of the stomach (middle) and bulged intestine (right) show alternating epithelial morphology and Ci-VSP expression. (D – F) Expression profile Ci-Dab2. (D) Whole-mount (left) 2-week-old juvenile showing regional expressions in the stomach (st; arrow), bulged-region of the mid-intestine (int; arrowhead), and posterior-intestine (gray arrowhead). Higher magnification (right) reveals a stripe-like distribution of signals in the stomach and bulged intestine. (E) Dissected post-pharyngeal region of the digestive tract of the 1.5-month-old young adult showing an additional expression region (white arrowhead) in the looped-intestine. (F) Transverse section specimens of the juvenile (white line in D) showing Ci-Dab2 expressions in the epithelium of the digestive tract (left). Expression signals in the stomach epithelium are restricted mainly in the inner-folds (white dotted lines) as opposed to the outer-folds (dotted lines). Circumferences of the stomach (middle) and bulged intestine (right) show alternating epithelial morphology and Ci-Dab2 expression. Scale bars = 200 µm in (A, D), 1 mm in (B, E), and 50 µm in (C, F). Additional abbreviations: os, oral siphon; as, atrial siphon; ph, pharynx; en, endostyle; stg, stigma; es, esophagus; nc, neural complex; gd, gonad.

Transverse sections of the digestive tract obtained from *Ciona* juveniles (Figure 7C) showed that the expression signals were present alternately in the digestive epithelial cells. In the stomach, Ci-VSP signals were observed mainly in the inner folds, where the epithelial cells predominantly express absorptive genes (Hayashibe et al., 2017). In the outer-folds of the stomach, known to express many pancreatic-related exocrine genes (Nakayama and Ogasawara, 2017), Ci-VSP signals were occasionally found in the regions which seemed to start invagination of newly generated inner-folds. Similarly, in the bulged region of mid-intestine, circumferential epithelial cells expressed Ci-VSP in a pattern corresponding to the alternating folding morphology as present in the stomach.

We further examined whether Ci-VSP expression correlates with endocytosis-associated genes. Several genes involved in enteric endocytosis in the zebrafish model have previously been identified, including Cubilin (*cubn*), Amnionless (*amn*), and Dab2 (*dab2*) (Park et al., 2019). In *Ciona*, however, only *dab2* ortholog (Ci-Dab2) is expressed as a single homolog, so Ci-Dab2 was selected to represent the endocytosis-associated gene in this study. In WISH on 2-week-old *Ciona* juveniles, the Ci-Dab2 expression pattern is highly comparable to that of Ci-VSP (Figure 7, D – F), implying their concerted function in the *Ciona* digestive tract. Ci-VSP is co-expressed with Ci-Dab2 in a strip-like pattern in the stomach and bulged intestine, indicating that Ci-VSP is expressed in the epithelial cells with endocytosis-dependent absorptive function.

## Discussion

VSP expression was previously reported in the digestive organs of other animal species, such as in the gut epithelial cells of invertebrate *C. intestinalis* Type A (Ogasawara et al., 2011) and the stomach of chicken embryos (Yamaguchi et al., 2014a). However, the functional role of VSP in the enterocytes has not been studied in detail. We report here the expression of VSP in specialized enterocytes known as LREs in the mid-intestine of zebrafish. VSP in LREs is most likely expressed on the endosomal membranes required for endocytosis. Loss of VSP activity impaired LRE endocytosis and nutrient absorption, potentially accounting for growth restriction and increased mortality in developing zebrafish. Dr-VSP was detectable in LREs as early as 5 dpf (Figure 1 and 2), coinciding with the time when zebrafish intestine begins nutrient absorption (Salinas and Parra, 2015; Lu et al., 2017). These findings highlight an essential physiological role of VSP in endocytosis-dependent nutrient absorption besides its previously established role in sperm physiology (Kawai et al., 2019).

### Localization of VSP on endomembranes

Previous research indicated that VSP is expressed on the plasma membrane, based on studies of native expression in *Ciona* sperms (Murata et al., 2005), *Xenopus* renal epithelial cells (Neuhaus and Hollemann, 2009; Ratzan et al., 2019), and mouse sperms (Kawai et al., 2019), as well as in vitro experiments using heterologous expression system (Murata et al., 2005; Ratzan et al., 2011; Mutua et al., 2014; Yamaguchi et al., 2014b). The same was reported for Dr-VSP when it was first characterized by whole-cell patch-clamp recordings of mammalian cells heterologously expressing Dr-VSP protein (Hossain et al., 2008). However, our results of immunofluorescence staining and immuno-electron microscopy revealed that native Dr-VSP in LREs was localized to the subapical region but not on microvilli nor plasma membrane (Figure 3 and Figure 4A). Results from the MDCK model indicate that Dr-VSP is enriched at the early and recycling endosomal membranes (Figure 4B), consistent with our recent in vitro study in which fluorescent signal from mCherry-fused Dr-VSP expressed in HEK293T cells was also distributed throughout the intracellular compartments (Kawanabe et al., 2020). Although additional studies, including in vivo co-localization analysis with endosome markers, are required, our current findings provide converging evidence that Dr-VSP functions primarily on endomembranes at the subapical region of LREs.

PIP conversion has been well documented to regulate membrane trafficking in the endosomal system, and this process is achieved by PIP-metabolizing enzymes, including PIP kinases and phosphatases (Di Paolo and De Camilli, 2006; Li et al., 2013; Wallroth and Haucke, 2018). Dr-VSP functions as 5-phosphatase and 3-phosphatase, allowing it to hydrolyze PI(3,4,5)P_3_, PI(4,5)P_2_, and PI(3,4)P_2_ (Keum et al., 2016; Okamura et al., 2018).

Following gavage with fDex and mCherry, we observed that endocytosis was remarkably decreased in Dr-VSP^-/-^ LREs, as evidenced by the reduction of subsequent endosomal vacuoles and fluorescence uptake into the cells (Figure 6). Histological and TEM findings show that loss of Dr-VSP resulted in fewer and smaller vacuoles and, consequently, a reduction in the overall cell volume of LREs. Besides, Dr-VSP^-/-^ LREs show reduced numbers of endocytic vesicles, cytoplasmic tubules, and tubule-vacuole complexes at the subapical region (Figure 7, Supplementary Figure S5 and Supplementary Figure S6). These findings suggest that Dr-VSP is involved in endocytic regulation, and Dr-VSP deficiency disrupts membrane trafficking in LREs.

### PIPs enzyme activity of VSP in endomembranes of LREs

Dr-VSP deficiency may alter PIP homeostasis on various types of endomembranes, which play a critical role in membrane trafficking. However, it remains a challenge to determine which enzyme subreaction mediated by VSP contributes to the phenotypes in LREs. Here we propose potential scenarios based on Dr-VSP’s dual phosphatase activities. One scenario considers a more robust Dr-VSP’s 5-phosphatase activity toward PI(4,5)P_2_ and PI(3,4,5)P_3_ (Keum et al., 2016). We speculate that intracellular Dr-VSP dephosphorylates such PIPs on the membranes of newly enclosed endocytic vesicles during the transformation to early endosomes (Zoncu et al., 2009; Vicinanza et al., 2011; Wang et al., 2019). Defective endocytosis in Dr-VSP^-/-^ LREs is likely caused by reduced dephosphorylation of endosomal PI(4,5)P_2_ and possibly PI(3,4,5)P_3_, leading to ectopic accumulation of these PIPs that prevent apical endocytic vesicles from transforming into, or fusing with, subsequent intracellular compartments during endosomal maturation; thus restricting most fDex and mCherry to the subapical region of Dr-VSP^-/-^ LREs. It should be noted that other 5-phosphatase, such as OCRL1 and INPP5B, can be detected in LREs, although at lower levels according to RNA-seq data (Park et al., 2019). These enzymes are involved in early endocytic trafficking (Erdmann et al., 2007; Vicinanza et al., 2011; Billcliff and Lowe, 2014; Cauvin et al., 2016) and can function similarly as VSP during active endocytosis. The presence of compensatory enzymes may account for the phenotypes of Dr-VSP^-/-^ LREs (Figure 6, F and H); some Dr-VSP^-/-^ zebrafish larvae survived and thrived into adulthood despite seemingly suffering from nutrient malabsorption.

Another scenario considers Dr-VSP’s 3-phosphatase activity towards PI(3,4)P_2_ and PI(3,4,5)P_3_ (Keum et al., 2016). Dr-VSP at recycling endosomes may be involved in the conversion of PI(3)P to PI(4)P by dephosphorylating the intermediary PI(3,4)P_2_ (Wang et al., 2019). Unless a compensatory mechanism is available, the loss of Dr-VSP’s 3-phosphatase activity can cause excessive PI(3,4)P_2_ retention and, as a result, disrupt the formation of recycling endosomes during endocytic recycling in LREs. The defect may also reduce subsequent endocytosis when the internalized membrane and specific receptors are unable to fully return to the plasma membrane. Furthermore, based on a previous study showing that VSP exhibits small but significant activity toward PI(3,5)P_2_ by in vitro enzyme assay (Matsuda et al., 2011), Dr-VSP may contribute to PI(3,5)P_2_ homeostasis in the late endocytic pathway in LREs (Wallroth and Haucke, 2018; Wang et al., 2019). The precise mechanism by which Dr-VSP is recruited to function at specific endomembranes has yet to be determined, and this will be one of the important issues to be addressed in the future for better understanding the molecular mechanisms underlying the physiological role of VSP in LREs.

Two distinct endocytic pathways have been described in zebrafish LREs: (i) fluid-phase endocytosis, for fDex uptake; and (ii) receptor-mediated endocytosis, for mCherry uptake (Park et al., 2019). In our study, Dr-VSP deficiency in zebrafish LREs affected fDex uptake more remarkably than mCherry uptake. Furthermore, following the gavage with fDex-mCherry mixture, the difference in the intracellular distribution of fDex and mCherry in LREs was more pronounced in Dr-VSP^-/-^ mutants (Supplementary Figure S4). These results suggest that the extent to which Dr-VSP contributes to LRE endocytosis differs between fluid-phase and receptor-mediated endocytosis, possibly due to subtle differences in molecular mechanisms, such as PIP conversion, between the two pathways.

### Roles of VSP in other tissues in zebrafish

Our immunostaining results indicate that Dr-VSP was also expressed in other tissues than the intestine. Among them, the expression in testis and kidney is consistent with the results obtained from other species (Tapparel et al., 2003; Neuhaus and Hollemann, 2009; Kawai et al., 2019; Ratzan et al., 2019), indicating that VSP is functionally conserved across animal species. We suspected that defective endocytosis in Dr-VSP^-/-^ LREs is most responsible for growth restriction and higher mortality in zebrafish larvae (Figure 5), particularly during the critical period when zebrafish larvae rely primarily on exogenous feeding via intestinal absorption (Ng et al., 2005; Wallace et al., 2005). Although Dr-VSP is also expressed in larval pronephros (Figure 2A), Dr-VSP deficiency in this organ is less likely to be the primary cause of mortality. This is because larval pronephros begins to function at around 48 hpf (Gerlach and Wingert, 2013), and defective pronephros should exhibit phenotypes earlier than what was observed in our study. We have previously reported that VSP regulates mouse sperm motility via its 5-phosphatase activity toward PI(4,5)P_2_ on the plasma membrane during sperm capacitation (Kawai et al., 2019). VSP deficiency in mice caused abnormal sperm movement and significantly reduced in vitro fertilization. Dr-VSP is expressed in zebrafish testis (Figure 1A) (Okamura et al., 2018), but Dr-VSP^-/-^ zebrafish have no apparent fertility defect. Further research may be necessary to determine whether Dr-VSP in zebrafish sperms serves a similar function as its ortholog in mouse sperms.

### Stimulus to activate VSP

VSP is a membrane protein that functions upon membrane depolarization (Murata et al., 2005; Okamura et al., 2009; Okamura et al., 2018). Our study raises an open question of how Dr-VSP is activated in LREs. According to the in vitro experiment, the voltage required to activate Dr-VSP is beyond the physiological range of membrane potential in native cells (Hossain et al., 2008). Unfortunately, there is no current in vivo method for measuring the membrane potential of intracellular vesicles in zebrafish LREs. However, several possible mechanisms may induce Dr-VSP activation in endosomes. First, unlike in vitro conditions, native Dr-VSP in LREs may function at physiological membrane potentials, possibly through interactions with unknown endogenous auxiliary subunits, and a partial activation could be sufficient to induce Dr-VSP phosphatase activity during active endocytosis. This is similar to what may happen in mouse sperms, where physiological membrane potential appears to activate mouse VSP (Kawai et al., 2019). Second, membrane dissociation during endosomal vesicle formation could alter membrane potential probably by changing membrane capacitance, ultimately accelerating depolarization. Third, changes in micro-environments such as membrane tension, luminal pH, and opening of endomembrane ion channels (Cang et al., 2014; Li et al., 2019; Wie et al., 2021) may facilitate Dr-VSP activation under physiological conditions. Further in vivo analyses are required to explore the relationship among membrane potential, VSP activation, and endosomal membrane trafficking. Potential future tests include rescue experiments of Dr-VSP^-/-^ mutants by expressing modified versions of VSP, such as a voltage-insensitive or a voltage-hypersensitive version, to study how membrane potential affects endocytosis in LREs. Additionally, optogenetics could be applied in zebrafish to manipulate membrane potential by specifically targeting photoactivated ion transporting molecules to specific membrane vesicles.

### Role of VSP shared among chordates

Our comparative study reveals that VSP is conserved in the digestive tracts of zebrafish and *C. intestinalis* Type A, as are other endocytosis-associated genes (Figure 2E and Figure 8, D – F) (Mishra et al., 2002; Maurer and Cooper, 2005; Park et al., 2019), suggesting that Dr-VSP and Ci-VSP share a conserved endocytic function in the absorptive epithelial cells. Endocytosis-dependent nutrient absorption is conserved across species from unicellular organisms to vertebrates. Unicellular organisms acquire nutrients from their surroundings by endocytosis, and many invertebrates show LRE-like gastrodermal cells that line the digestive tract to absorb nutrients by endocytosis (Hartenstein and Martinez, 2019). Endocytotic absorption through epithelial cells has also been reported in the nutrient absorptive organs, such as the trophotaenia of viviparous fish embryos (Iida et al., 2021) and the ileum of pre-weaning mammals (Kraehenbuhl and Campiche, 1969; Gonnella and Neutra, 1984). Furthermore, LREs have also been proposed to be involved in trans-cellular transport of antibodies and other antigens across enterocytes to extra-intestinal tissues for immune modulation (Kraehenbuhl and Campiche, 1969; Kleinman and Walker, 1984; Wallace et al., 2005; Brugman, 2016; Park et al., 2019). Because both of LRE-like cells and VSP gene are widely observed from Cnidaria to mammals (Kraehenbuhl and Campiche, 1969; Gonnella and Neutra, 1984; Lickwar et al., 2017; Hartenstein and Martinez, 2019; Park et al., 2019), the endocytic role of VSP may well be conserved among a wide range of animal species.

## Materials and Methods

### Experimental model and subject details

#### Zebrafish maintenance

We used RIKEN wild-type zebrafish (*Danio rerio*) obtained from RIKEN Brain Science Institute (Saitama, Japan), and *Tg(vsp:EGFP)* transgenic zebrafish that express EGFP under the endogenous expression of *vsp*, the Dr-VSP-encoding gene. The fish were fed a non-calorie-restricted standard diet twice daily and raised in an aquatic system maintained at 28°C with a 14 h/10 h light/dark cycle. All experiments were conducted under the regulation of the Animal Care Facility, Center for Medical and Translational Research (CoMIT) at Osaka University (Osaka, Japan).

#### Biological materials for Ciona intestinalis’s experiments

Matured and young adult *Ciona intestinalis* type A (*C. robusta*) were collected during the spawning season at Port Chiba in Tokyo Bay (Chiba, Japan). *Ciona* Juveniles were developed by artificial insemination using matured eggs and sperms obtained from dissected gonoduct. The specimens of whole-mount juveniles and dissected post-pharyngeal region of the digestive tracts of the young adult were prepared for in situ hybridization as previously described (Nakayama and Ogasawara, 2017).

### Method details

#### RNA isolation and Reverse Transcription PCR (RT-PCR)

Total RNA was purified from zebrafish larvae and adult tissues using TRIzol™ LS reagent (Invitrogen, USA) with volume adjustment for small samples. Total RNA (200 ng) extracted from each sample was reverse-transcribed using SuperScript™ III First-Strand Synthesis System (Invitrogen, USA). For semi-quantitative RT-PCR, fragments of *vsp* and β*- actin* were amplified from cDNA using the primer sets listed in Supplementary Table S1.

#### Whole-mount in situ hybridization (WISH) for zebrafish larvae

Zebrafish larvae were fixed in 4% PFA/PBS at 4ºC overnight. Next, the larvae were washed twice with PBS for 5 min each time, followed by briefly washed with 100% MeOH and kept in MeOH at -20ºC until the following steps. WISH was performed using a digoxigenin (DIG)-labeled RNA probe according to the standard protocol. In this study, the Dr-VSP cDNA was subcloned into pBluescript II KS (+) plasmid vector. The resultant plasmid was linearized by XhoI enzyme and later used as a template for synthesizing DIG-labeled RNA probe using T3 RNA polymerase. At the beginning of WISH process, fixed zebrafish larvae were treated with proteinase K (10µg/mL) for 15 min at room temperature. Prehybridization was done at 55ºC for 1 h. Larva samples were hybridized with 10 ng of DIG-labeled Dr-VSP cRNA probe overnight. During posthybidization, larva samples were washed with PBS containing 0.2% Tween-20 and then incubated with anti-DIG-alkaline phosphatase (AP), Fab fragments (Roche, Mannheim, Germany) at 4ºC overnight. The samples were subsequently washed with PBS containing 0.2% Tween-20 every 20 min for 3 h, to remove excessive antibody. Positive signals of the hybridized Dr-VSP mRNA transcript were visualized by color development using BCIP/NBT solution (Wako, Japan) for 2 h at room temperature. The stained larvae were observed under Zeiss Stemi SV11 Stereo Microscope (Zeiss, Berlin, Germany).

#### In situ hybridization, detection, and observation of gene expression in Ciona intestinalis

DIG-labeled antisense RNA probe for *Ciona* genes were synthesized from T7 RNA polymerase promoter-attached amplified cDNA, and probes were purified by centrifugal ultrafilter as previously described (Ogasawara et al., 2011). WISH of the *Ciona* specimens was performed using “InSitu Chip” (Ogasawara et al., 2006). Gene expression was visualized using BCIP/NBT solution (Roche, Mannheim, Germany). After color development, the digestive tracts of the *Ciona* juveniles were sectioned transversely with a thickness of 10 μm (Nakayama and Ogasawara, 2017). Whole-mount and sectioned specimens of *Ciona* were observed under BX51 microscope (Olympus, Japan) or SZX12 Stereo Microscope (Olympus, Japan).

#### CRISPR-Cas9-mediated transgenesis and mutagenesis

*Tg(vsp:EGFP)* zebrafish line was generated via CRISPR-Cas9-mediated knock-in transgenesis (Kimura et al., 2014) with adaptation to the *vsp* gene (Supplementary Figure S1). The sgRNA for genome digestion contained a target site (AAACCGCTACGTCTGCAGCAGG) derived from the *vsp* exon sequence located between the transcriptional start site and translational start site. Donor DNA for knock-in contained Mbait-hsp70 promoter-EGFP-polyA sequences. Successful knock-in transgenesis was validated by sequence mapping for donor DNA insertion and comparing EGFP signal with *vsp* expression pattern. Dr-VSP-deficient (Dr-VSP^-/-^) zebrafish were generated via CRISPR-Cas9-mediated mutagenesis. Potential CRISPR targets for the *vsp* gene were identified via the online tool CHOPCHOP (Montague et al., 2014) based on genomic DNA sequence obtained from the Ensembl zebrafish genome database version 9 (Zv9, https://asia.ensembl.org/Danio_rerio/). We selected three targets close to the translation initiation site without potential off-target (Supplementary Table S2). In our setting, target T1 in exon 6 (Supplementary Figure S3) yielded the highest efficiency in producing adequate frameshift mutation. Microinjection for Dr-VSP mutagenesis was performed base on the reported protocol (Kotani et al., 2015). All injected embryos were incubated at 28.5°C to support early development. Subsequently, some injected embryos underwent genome extraction and sequence analysis, while the remaining embryos were raised as the founder (F0) generation.

#### Genomic DNA extraction and sequence analysis

Genomic DNA was isolated from whole larvae or fin-clips of adult zebrafish using the HotSHOT DNA extraction procedure (Meeker et al., 2007). Genomic DNA fragments covering CRISPR targets were amplified by PCR using locus-specific primer sets listed in Supplementary Table S2. We used BigDye™ Terminator v3.1 Cycle Sequencing Kit (Thermo Fisher Scientific, Waltham, MA) for analyzing the sequences of PCR fragments.

#### Fish sectioning, immunohistochemistry (IHC), and toluidine staining

Zebrafish larvae were fixed in 4%PFA/PBS, followed by 30% sucrose/PBS, and embedded in 5% ultra-low gelling temperature agarose (Sigma-Aldrich, USA). Embedded specimens were cryosectioned in 10 µm slices using Leica CM3050S cryostat (Leica, Germany). Antibodies used for IHC are as follows: anti-Dr-VSP/TPTE (NeuroMab, USA), Acti-stain™ 488 Phalloidin (Cytoskeleton, USA), Alexa Fluor™ 594 Phalloidin (Invitrogen, USA), and anti-Na^+^-K^+^ ATPase (Ab76020; Abcam, USA). Images were taken under Zeiss LSM880 confocal microscope with Airyscan (Zeiss, Berlin, Germany). For toluidine blue staining, larvae were fixed in 4%PFA/PBS at 4°C overnight, followed by 2.5%GA at 4°C for 2 h and 1%GA overnight. Samples were subsequently dehydrated by ethanol substitution and embedded in Epon for thin sectioning before the staining process.

#### Transmission electron microscopy (TEM)

Zebrafish were fixed in 4%PFA at 4°C overnight before en-bloc sectioning of the mid-intestine region. Sectioned samples were subsequently fixed in 2.5%GA at 4°C for 2 h, followed by 1%GA overnight. Specimen preparation and staining were done as standard protocol, with technical support from Center for Medical Research and Education, Osaka University. Briefly, the fixed samples were incubated with 1% osmium tetroxide, then dehydrated in ethanol series and propylene oxide. The samples were then embedded in Epon and ultra-thin sectioned, followed by contrast stained with uranyl acetate and lead citrate. TEM images were acquired under the Hitachi H-7650 transmission electron microscope (Tokyo, Japan) operating at 80 kV, and analyzed using ImageJ software (NIH, USA).

#### Pre-embedding immuno-electron microscopy (Immuno-EM)

Zebrafish larvae were fixed in 4%PFA/HEPES, or 4%PFA/0.1%GA/HEPES, before being embedded in 5% ultra-low gelling temperature agarose (Sigma-Aldrich, USA) and cryosectioned in 10 µm slices using Leica CM3050S cryostat (Leica, Germany). Sectioned samples were incubated in 5% normal goat serum (NGS)/0.125% saponin/PBS for 30 min, then incubated in anti-Dr-VSP/TPTE antibody (clone N432/21, NeuroMab, USA) at 37°C for 2 h, followed by Alexa Fluor® 488 FluoroNanogold antibody (Nanoprobes, USA) at room temperature for 2 h. Immunogold particles were intensified at room temperature with HQ SILVER™ enhancement kit (Nanoprobes, USA) and 5% selenium toner. Samples were further treated with 1% osmium tetroxide, 0.5% uranyl acetate, and underwent the same process as described in the TEM protocol.

#### Cell culture and transfection

MDCK type 2 (MDCKII) cells (CRL-2936, ATCC) were cultured in D-MEM high glucose (Wako, Japan) containing 10% FBS at 37ºC under 5% CO_2_. Transient expression was achieved by co-transfecting mCherry-Dr-VSP plasmid with (i) EGFP-Rab5 plasmid (early endosome marker) or (ii) EGFP-Rab11 plasmid (recycling endosome marker) into MDCKII cells using Lipofectamine™ 3000 transfection reagent (Invitrogen, USA).

#### Fish gavage and live imaging

6-dpf zebrafish larvae were gavaged with Alexa Fluor 488-tagged dextran 10,000MW (fDex) solution (Molecular Probes, USA) or purified mCherry solution (prepared in the laboratory) following the established method (Cocchiaro and Rawls, 2013) (see also Figure 6A). A microforged capillary needle loaded with either solution was directly inserted through the larval mouth to inject the solution into the anterior intestinal bulb. Gavaged larvae were incubated at 28.5°C for 2 h to allow intestinal absorption before live image acquisition under Zeiss 710 confocal microscope (Zeiss, Berlin, Germany) with temperature control at 28.5°C for entire sessions. The following parameters were analyzed: (i) the percentage of vacuolated cells per section; and (ii) relative fDex or mCherry internalization profile measured across the apicobasal axis of individual cells and the longitudinal axis along the posterior portion of mid-intestine. The data of (ii) were analyzed by measuring fluorescence intensity using Plot Profile function in ImageJ software (NIH, USA) as previously described (Park et al., 2019) and presented as means ± s.e.m. percentage of intracellular fluorescence intensity along the specified axes, with a 100% indicates the maximal value of the average intensity on each axis.

### Quantification and statistical analysis

Sample sizes, definitions of replicates, and statistical details for each experiment are reported in the figure legends. Data are presented as mean ± s.e.m. unless otherwise specified. P value < 0.05 was regarded as statistically significant. Graphs and statistical analyses were performed using Prism 8 (Graphpad Software, San Diego, CA).

#### Parameter analysis of larval enterocytes

Quantification of cellular dimensions of LREs was done based on the previous procedure (Sidhaye et al., 2016). We performed immunostaining with Acti-stain™ 488 Phalloidin (Cytoskeleton, USA) and anti-Na^+^-K^+^ ATPase Antibody (Ab76020; Abcam, USA) to outline individual cell (Figure 7A). In the mid-intestine region, each confocal image was selected where enterocytes appeared the broadest, and the following parameters were analyzed: central height, apical width, basal width, and microvillus length.

#### Body length and survival analysis

Wild-type and Dr-VSP^-/-^ zebrafish were raised in the standard system of the Animal Care Facility, Center for Medical and Translational Research (CoMIT), Osaka University. Body length and remaining number of larvae were measured periodically from the day of fertilization to 28 dpf. All data were collected from 3 sets of independent experiments.

#### Analysis of gene expression level using RNA-seq data

Raw data of the RNA-seq experiments were retrieved from the GEO Datasets database with accession number GSE124970 (Park et al., 2019). Heatmap representing expression levels of selected genes in IECs and LREs (Figure 2E) was analyzed using web-based application iDEP v0.92 (http://bioinformatics.sdstate.edu/idep92) (Ge et al., 2018)

#### Image processing

Raw images were processed and analyzed using ImageJ software (NIH, USA).

## Supporting information

DrVSP Supplementary Figures

DrVSP Supplementary Table S1

DrVSP Supplementary Table S2

DrVSP Supplementary Table S3

DrVSP Supplementary Movie S1

DrVSP Supplementary Movie S2

## Data availability

Data generated or analyzed in the present study are included in this published article and its supplementary information files. Key reagents and resources are listed in Supplementary Table S3. Further information and requests should be directed to and will be fulfilled by the corresponding author.

## Acknowledgments

We thank Dr. Shinji Kanda (University of Tokyo, Japan) for helpful advice on this study. Our thanks are also due to Drs. Junji Takeda, Akihiro Harada, Shin-ichiro Yoshimura, Atsushi Tamura, Hiroo Tanaka, Yoshifumi Okochi, Akira Kawanabe (Osaka University, Japan), and all laboratory members for valuable suggestions and discussions. We thank Mr. Eiji Oiki (Osaka University, Japan) for advice in electron microscopy; Mr. Masayuki Yamagishi and Mr. Satoshi Nakayama (Chiba University, Japan) for advice and technical assistance in the in-situ hybridization of the *Ciona* specimens. Technical support for this study was provided by the Center for Medical Research and Education and the Center for Medical and Translational Research (CoMIT) at Osaka University.

## Additional information

## Author contributions (CRediT taxonamy)

Conceptualization and Methodology, A.R., T.K., and Y.O.; Investigation, A.R., M.M., S.H., Y.K., F.T., T.M., M.I.H., and M.O.; Formal Analysis, A.R., T.K., and Y.O.; Visualization and Writing - Original Draft, A.R.; Writing - Review & Editing, A.R., T.K., Y.O.; Funding Acquisition, T.K., Y.O., A.R.; Supervision, Y.O.

## Ethics

Protocols used for the animal experiments in this study were approved by the Animal Research Committee of Osaka University, Japan (No. 27-079). All procedures were conducted in accordance with the regulation of the Animal Care Facility, Center for Medical and Translational Research (CoMIT) at Osaka University, Japan.

## Figure legends for supplementary figures

**Supplementary Figure S1: Experimental schemes of CRISPR-Cas9-mediated *Tg(vsp:EGFP)* transgenesis**

Generation of *Tg(vsp:EGFP)* transgenic zebrafish. Donor DNA plasmid containing the enhanced green fluorescence protein (EGFP) sequence was incorporated into *vsp* exon1, which is located between transcription start site (exon1) and translation start site (exon3). Successful transgenesis expresses EGFP recapitulating endogenous expression of *vsp* gene. For detailed information, please refer to the Method Details, and Kimura et al (Kimura et al., 2014). E1, exon 1. E2, exon2. hsp, hsp70 promoter. pA, polyA.

**Supplementary Figure S2: Spatial distribution of Dr-VSP within LREs.**

Representative images of pre-embedding Dr-VSP immunogold staining in LREs from 14-dpf wild-type zebrafish larva, demonstrating the spatial distribution of Dr-VSP signals at multiple areas within the cell. Dr-VSP is highly expressed at the subapical region (inset 1) and gradually decreased in the middle (inset 2) and basal part (inset 3) of LREs, corresponding to the immunostaining results in Figure 3. Scale bar = 2 µm. V, vacuole. N, nucleus.

**Supplementary Figure S3: Experimental schemes of CRISPR-Cas9-mediated Dr-VSP**^**-/-**^ **mutagenesis.**

(A – E) Generation of heritable Dr-VSP^-/-^ zebrafish. (A) Schematic diagram of the zebrafish Dr-VSP-encoding gene (*vsp*), consisting of 21 exons. The CRISPR target with the protospacer adjacent motif (PAM) sequence (TGG) is indicated in the box.

(B) Sequencing results illustrate microdeletion (arrowhead) in the genomic sequence of Dr-VSP^-/-^ zebrafish. WT, wild type.

(C) Alignment of genomic DNA sequences (top) and amino acid sequences (bottom) comparing between wild-type and Dr-VSP^-/-^ zebrafish. Microdeletion (red) within the CRISPR target (gray shade) induced frameshift mutation in the subsequent amino acids of Dr-VSP^-/-^ zebrafish.

(D) Model of Dr-VSP illustrates the CRISPR-Cas9-mediated mutagenesis in topological view. Mutation disrupted the functional domains of Dr-VSP located distal to the CRISPR target.

(E) Immunostaining of wild-type and Dr-VSP^-/-^ zebrafish. Anti-Dr-VSP antibody is specific to the S4 transmembrane region and intracellular active center of the phosphatase domain. Dr-VSP^+^ cell is undetectable in Dr-VSP^-/-^ enterocytes. Green, Dr-VSP. Red, F-actin. Blue, DAPI. Scale bar = 10 µm

**Supplementary Figure S4: Preliminary results of fDex-mCherry mixture internalization into zebrafish LREs after gavage.**

(A) Live confocal images of wild-type (WT) LREs showing fDex-mCherry mixture internalization. Both fDex and mCherry signals are located in apical vesicles and supranuclear vacuoles (arrowheads). Some mCherry signals did not overlap with fDex signals (as indicated by asterisks).

(B) Live confocal images of Dr-VSP^-/-^ LREs showing fDex-mCherry mixture internalization. Notably, mCherry was transported deeply into some supranuclear vacuoles (arrowheads), whereas fDex was mostly found in apical vesicles. Dotted lines indicate the outlines of larval intestines. Many mCherry signals did not overlap with fDex signals (as indicated by asterisks), indicating that mCherry and fDex were uptaken via different mechanisms into distinct intracellular vesicles. The mismatch between mCherry and fDex signals was more pronounced in Dr-VSP^-/-^ than wild-type LREs. Scale bar = 20 µm.

**Supplementary Figure S5: Morphological structures of zebrafish LREs under transmission electron microscopy.**

(A) Representative TEM images (left) of LREs from 14-dpf wild-type (top) and Dr-VSP^-/-^ (bottom) zebrafish larvae; and their corresponding schematic illustration (right). Green color represents the subapical region upper to supranuclear vacuoles, where the positive immunofluorescence signal of Dr-VSP was observed in Figure 3, A and B. N, nucleus. V, vacuole. Scale bar = 10 μm.

(B) Representative TEM images of LREs from 14-dpf wild-type (top) and Dr-VSP^-/-^ (bottom) zebrafish larvae, showing the ultrastructures of microvilli and subapical region. Key features of absorptive enterocytes are presented, including membrane invaginations at inter-microvillous spaces, cytoplasmic tubules and tubule-vacuole complexes, and numerous endocytic vesicles. In wild-type LRE, green color represents the subapical region upper to supranuclear vacuole, where immunofluorescence signal of Dr-VSP was observed in Figure 3, A and B. Scale bar = 2 μm (left) and 500 nm (right).

(C) Representative TEM images of LREs from 14-dpf wild-type (left) and Dr-VSP^-/-^ (right) zebrafish larvae, showing the ultrastructures of microvilli and subapical region. Red color represents the area beneath apical surface that contains cytoplasmic tubules but few well-defined endocytic vesicles. The vesicles are more densely distributed in the areas deeper than the red region and are frequently associated with larger endosomal vacuoles.

(D) Inter-edge distance between apical surface and vesicle-dense area in wild-type and Dr-VSP^-/-^ LREs, corresponding to the width of red shading areas shown in (C). Enterocytes, > 50 cells from 25 TEM images (magnification = 15000x) for each zebrafish line. Data were collected from 4 wild-types and 3 Dr-VSP^-/-^ larvae. Average distances in each TEM image were analyzed using ImageJ macro.

(E) Microvillous length of wild-type and Dr-VSP^-/^ LREs measuring at TEM level from the same samples as in (D). Error bars, means ± s.d.; ****P < 0.0001; unpaired Student’s t-test; ns, no statistically significant difference.

**Supplementary Figure S6: TEM image series of wild-type and Dr-VSP**^**-/-**^ **LREs.**

Representative TEM image series of LREs from 14-dpf wild-type (top) and Dr-VSP^-/-^ (bottom) zebrafish larvae. Large vacuoles are distributed in the supranuclear region but are fewer in Dr-VSP^-/-^ LREs. Membrane invaginations at inter-microvillous spaces (*) can be observed at higher magnification. Branching cytoplasmic tubules and tubule-vacuole complexes (arrowheads) are more common in wild-type LREs. Scale bar at 6000x = 10 μm; at 15000x = 2 μm; at 40000x = 500 nm.

**Supplementary Figure S7:**

Multiple amino acid alignment of full-length VSP orthologs from human (*Homo sapiens*, Hs-VSP), mouse (*Mus musculus*, Mm-VSP), zebrafish (*Danio rerio*, Dr-VSP), common carp (*Cyprinus carpio*, Cc-VSP), Nile tilapia (*Oreochromis niloticus*, On-VSP), Japanese medaka (*Oryzias latipes*, Ol-VSP), sea squirt (*Ciona intestinalis* Type A, Ci-VSP), and Crown-of-thorns starfish (*Acanthaster planci*, Ap-VSP). Boxes indicate the putative transmembrane helices (S1 - S4), VSD-PD linker, WPD loop, CX5R(T/S), gating loop, 515 loop/CBR3 loop (Okamura et al., 2018). Amino acid sequences were retrieved from National Center for Biotechnology Information (NCBI) public database. Analyses were performed using online Constraint-based Multiple Alignment Tool (COBALT) (https://www.ncbi.nlm.nih.gov/tools/cobalt/)

**Supplementary Figure S8:**

Molecular phylogenetic tree showing the phylogenic relationships among VSP orthologs from **mammals:** human (*Homo sapiens*, Hs-VSP1 and Hs-VSP2), mouse (*Mus musculus*, Mm-VSP), rat (*Rattus norvegicus*, Rn-VSP), dog (*Canis lupus familiaris*, Clf-VSP), cat (*Felis catus*, Fc-VSP); **amphibian:** American clawed frog (*Xenopus laevis*, Xl-VSP1 and Xl-VSP2); **teleosts:** goldfish (*Carassius auratus*, Ca-VSP), common carp (*Cyprinus carpio*, Cc-VSP), zebrafish (*Danio rerio*, Dr-VSP), channel catfish (*Ictalurus punctatus*, Ip-VSP), Chinook salmon (*Oncorhynchus tshawytscha*, Ots-VSP), Atlantic salmon (*Salmo salar*, Ss-VSP), Siamese fighting fish (*Betta splendens*, Bs-VSP), Japanese puffer (*Takifugu rubripes*, Tr-VSP), turquoise killifish (*Nothobranchius furzeri*, Nf-VSP), Japanese medaka (*Oryzias latipes*, Ol-VSP), Nile tilapia (*Oreochromis niloticus*, On-VSP); **invertebrates:** sea squirt (*Ciona intestinalis* Type A, Ci-VSP), crown-of-thorns starfish (*Acanthaster planci*, Ap-VSP), fresh-water polyp (*Hydra vulgaris*, Hv-VSP), sea anemone (*Exaiptasia diaphana*, Ed-VSP), and smooth cauliflower coral (*Stylophora pistillata*, Spi-VSP). Data were retrieved from the same source as Supplementary Figure S7, and analyzed using Molecular Evolutionary Genetics Analysis (MEGA) X Software (Pennsylvania State University, Pennsylvania, US).

**Supplementary Figure S9: Morphological structure of ascidian intestine and expression profiles of intestine-related genes**

(A – D) Expression profiles of intestine-related genes in *Ciona* juvenile (1^st^ column) and young adult (2^nd^ column) revealed by WISH. In the juvenile, whole (left panel) and magnificent of the post-pharyngeal region of the digestive tract including esophagus, stomach, and intestine (right panel) were shown.

(A) Morphological structure of the digestive tract was noted on the expressions of Mucin gene. Digestive tract of *C. intestinalis* Type A was divided into oral siphon (os), pharynx (ph), esophagus (es), stomach (st), and intestine (int) (Chiba et al., 2004). The intestine was subdivided into four regions, R1 to R4 (Yoshida and Sasakura, 2012). Mucin gene was expressed broadly in the post-pharyngeal region of the digestive tract of the juvenile, but not expressed in the stomach and bulged-intestine (R3). In the young adult specimen, expression profile was similar to the juvenile, but expression signal was disappeared between anterior intestine (R1) and looped-intestine (R2) beside the developing gonad (gd).

(B) Absorption-related PEPT1 gene was invested as an example of the peptide transporter genes. Expression signals of the *PEPT1* were observed in stomach (arrow) and bulged-intestine (R3) of the mid-intestine (arrowhead) in both developmental stages of juvenile and young adult. Additional expression (white arrowhead) was appeared between anterior intestine (R1) and looped-intestine (R2) of the young adult intestine.

(C) Absorption-related SGLT1 gene was invested as a secondary active glucose transporter. Transcripts of the *SGLT1* were detected in stomach (arrow), bulged-intestine (R3) (arrowhead), and posterior-intestine (R4) (gray arrowhead) in juvenile and young adult specimens.

(D) GLUT5 gene was also assess as a facilitated transporter of the monosaccharide. Transcripts of the *GLUT5* were detected in stomach (arrow), bulged-intestine (R3) (arrowhead), and posterior-intestine (R4) (gray arrowhead) in juvenile and young adult specimens. Additional expression (white arrowhead) between anterior intestine (R1) and looped-intestine (R2) was also found in the young adult. Scale bars = 200 µm in (left) and 1 mm in (right). Additional abbreviations: as, atrial siphon; en, endostyle. Latest gene model IDs (HT version) of the *Ciona Mucin, PEPT1, SGLT1*, and *GLUT5* are KY.Chr2.2226, KY.Chr13.370, KY.Chr10.1303, and KY.Chr7.810, respectively (Satou et al., 2019)

**Supplementary Movie S1: Dr-VSP is localized at the endosomal membranes of early endosomes.**

MDCKII cells co-expressing Dr-VSP-mCherry and Rab5-EGFP for early endosomes. Time-lapse images were acquired from the same samples as presented in Figure 4B. Red, Dr-VSP. Green, Rab5. Scale bar = 20 μm.

**Supplementary Movie S2: Dr-VSP is localized at the endosomal membranes of recycling endosomes.**

MDCKII cells co-expressing Dr-VSP-mCherry and Rab11-EGFP for recycling endosomes. Dr-VSP is localized at the endosomal membranes of recycling endosomes. Time-lapse images were acquired from the same samples as presented in Figure 4B. Red, Dr-VSP. Green, Rab11. Scale bar = 20 μm.

**Supplementary Table S1: RT-PCR Primers used in this study**

**Supplementary Table S2: CRISPR target sequences of *vsp* and PCR primers used in this study**

**Supplementary Table S3: Key resources table**

