## Supplementary material for "Voltage-Sensing Phosphatase (VSP) Regulates Endocytosis-Dependent Nutrient Absorption in Chordate Enterocytes": DrVSP Supplementary Figures

### Supplementary Figure S1

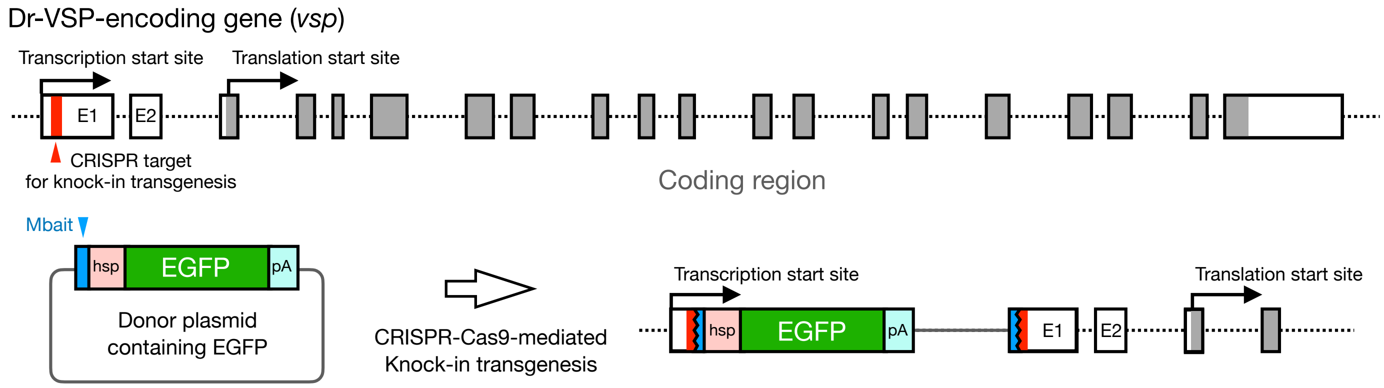

#### Figure legends for supplementary figures

**Supplementary Figure S1: Experimental schemes of CRISPR-Cas9-mediated *Tg(vsp:EGFP)* transgenesis**

Generation of *Tg(vsp:EGFP)* transgenic zebrafish. Donor DNA plasmid containing the enhanced green fluorescence protein (EGFP) sequence was incorporated into *vsp* exon1, which is located between transcription start site (exon1) and translation start site (exon3). Successful transgenesis expresses EGFP recapitulating endogenous expression of *vsp* gene. For detailed information, please refer to the Method Details, and Kimura et al (Kimura et al., 2014). E1, exon 1. E2, exon2. hsp, hsp70 promoter. pA, polyA.

### Supplementary Figure S2

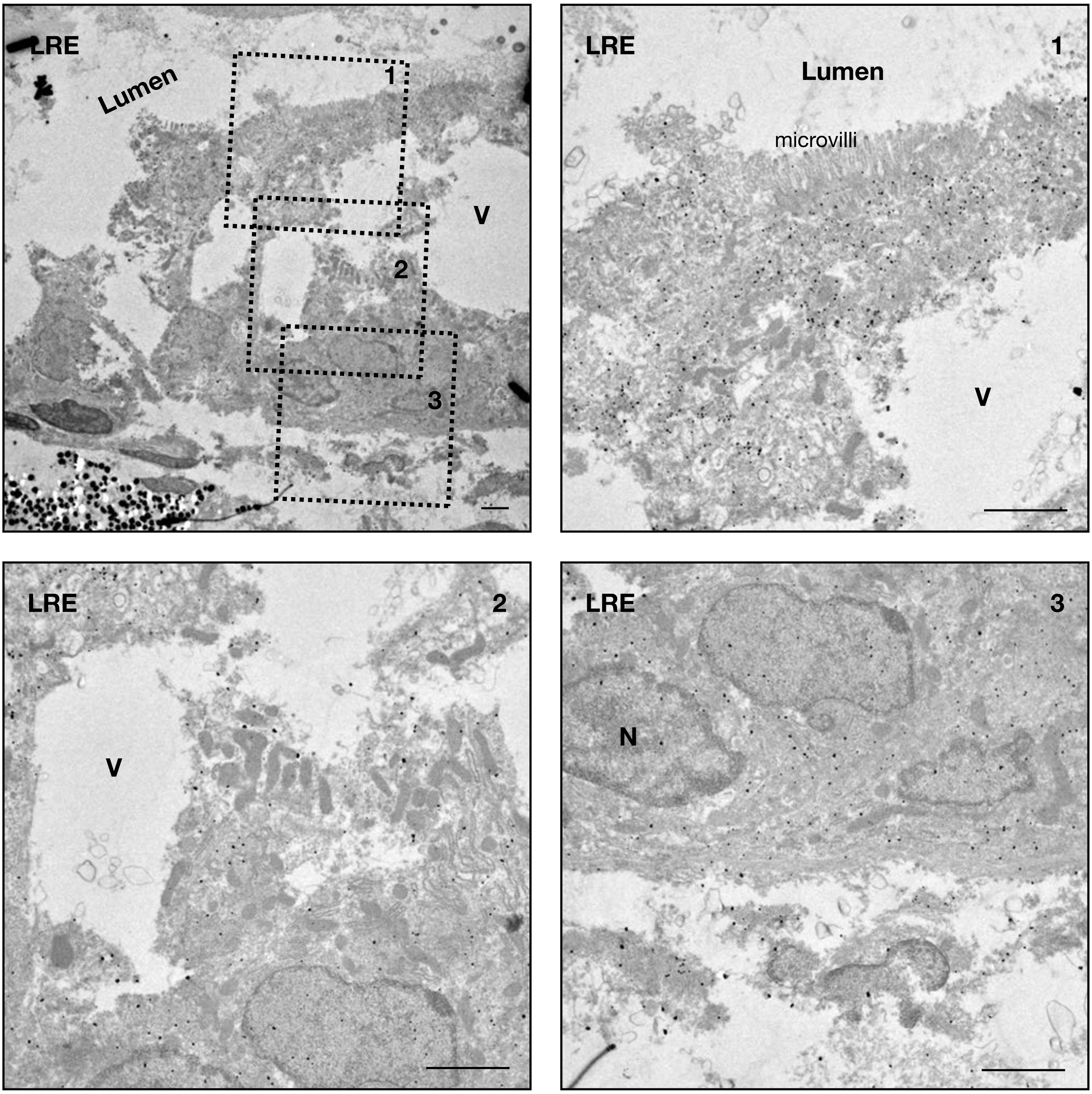

**Supplementary Figure S2: Spatial distribution of Dr-VSP within LREs.**

Representative images of pre-embedding Dr-VSP immunogold staining in LREs from 14-dpf wild-type zebrafish larva, demonstrating the spatial distribution of Dr-VSP signals at multiple areas within the cell. Dr-VSP is highly expressed at the subapical region (inset 1) and gradually decreased in the middle (inset 2) and basal part (inset 3) of LREs, corresponding to the immunostaining results in Figure 3. Scale bar = 2 μm. V, vacuole. N, nucleus.

### Supplementary Figure S3

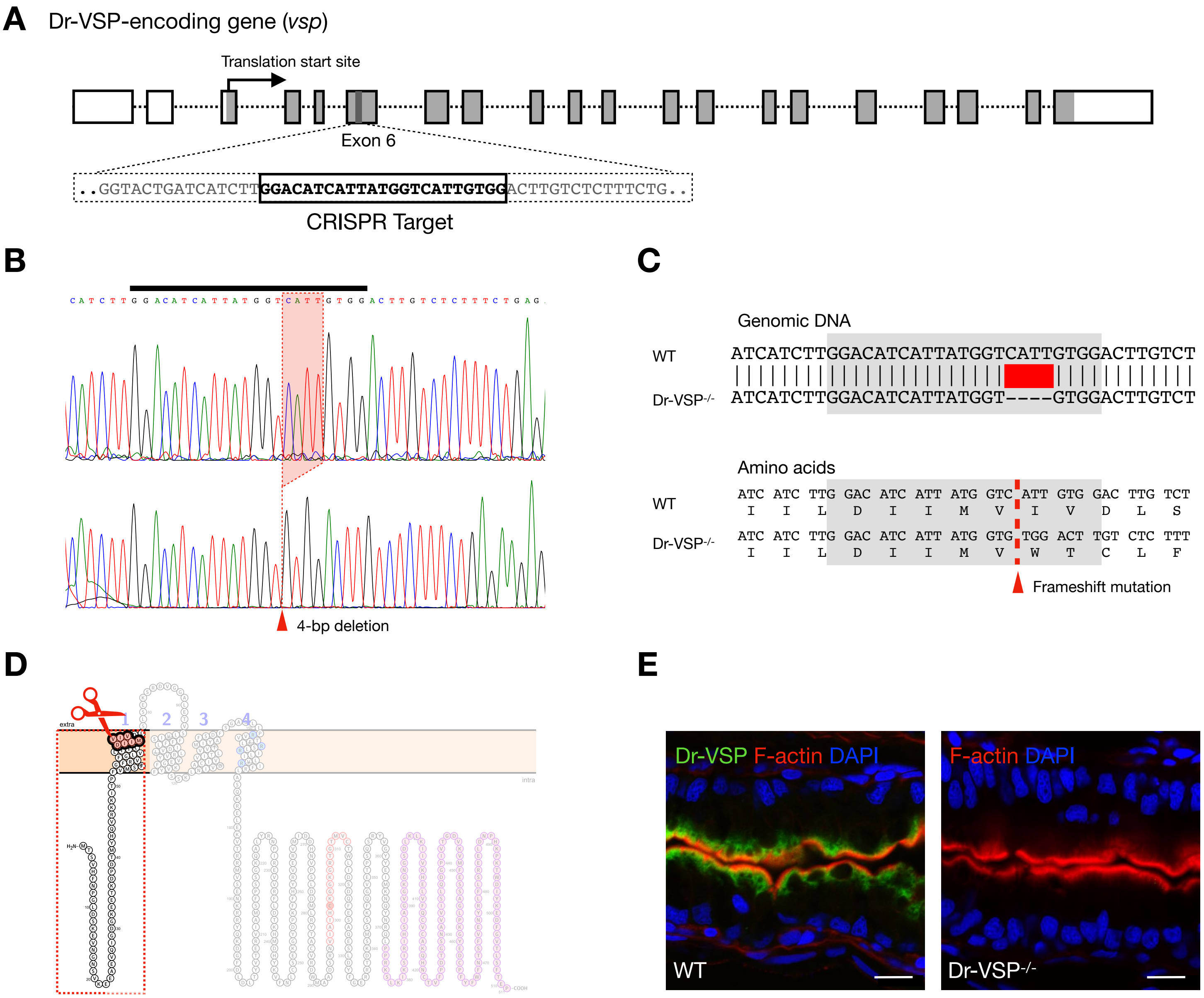

**Supplementary Figure S3: Experimental schemes of CRISPR-Cas9-mediated Dr-VSP$^{-/-}$ mutagenesis.**

(A – E) Generation of heritable Dr-VSP$^{-/-}$ zebrafish. (A) Schematic diagram of the zebrafish Dr-VSP-encoding gene (*vsp*), consisting of 21 exons. The CRISPR target with the protospacer adjacent motif (PAM) sequence (TGG) is indicated in the box.

(B) Sequencing results illustrate microdeletion (arrowhead) in the genomic sequence of Dr-VSP$^{-/-}$ zebrafish. WT, wild type.

(C) Alignment of genomic DNA sequences (top) and amino acid sequences (bottom) comparing between wild-type and Dr-VSP$^{-/-}$ zebrafish. Microdeletion (red) within the CRISPR target (gray shade) induced frameshift mutation in the subsequent amino acids of Dr-VSP$^{-/-}$ zebrafish.

(D) Model of Dr-VSP illustrates the CRISPR-Cas9-mediated mutagenesis in topological view. Mutation disrupted the functional domains of Dr-VSP located distal to the CRISPR target.

(E) Immunostaining of wild-type and Dr-VSP$^{-/-}$ zebrafish. Anti-Dr-VSP antibody is specific to the S4 transmembrane region and intracellular active center of the phosphatase domain. Dr-VSP$^{+}$ cell is undetectable in Dr-VSP$^{-/-}$ enterocytes. Green, Dr-VSP. Red, F-actin. Blue, DAPI. Scale bar = 10 µm

### Supplementary Figure S4

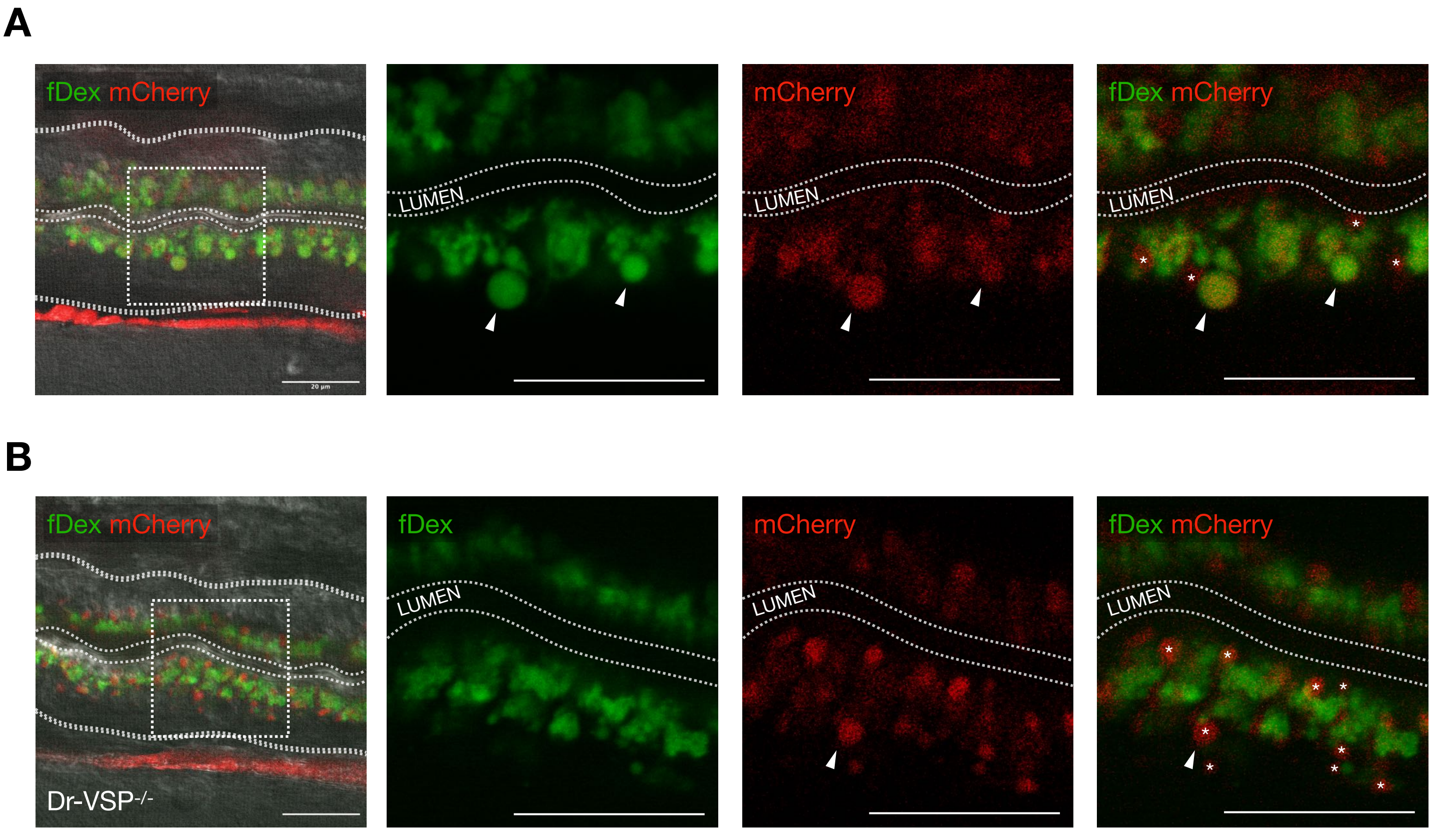

**Supplementary Figure S4: Preliminary results of fDex-mCherry mixture internalization into zebrafish LREs after gavage.**

(A) Live confocal images of wild-type (WT) LREs showing fDex-mCherry mixture internalization. Both fDex and mCherry signals are located in apical vesicles and supranuclear vacuoles (arrowheads). Some mCherry signals did not overlap with fDex signals (as indicated by asterisks).

(B) Live confocal images of Dr-VSP$^{-/-}$ LREs showing fDex-mCherry mixture internalization. Notably, mCherry was transported deeply into some supranuclear vacuoles (arrowheads), whereas fDex was mostly found in apical vesicles. Dotted lines indicate the outlines of larval intestines. Many mCherry signals did not overlap with fDex signals (as indicated by asterisks), indicating that mCherry and fDex were uptaken via different mechanisms into distinct intracellular vesicles. The mismatch between mCherry and fDex signals was more pronounced in Dr-VSP$^{-/-}$ than wild-type LREs. Scale bar = 20 μm.

#### Supplementary Figure S5

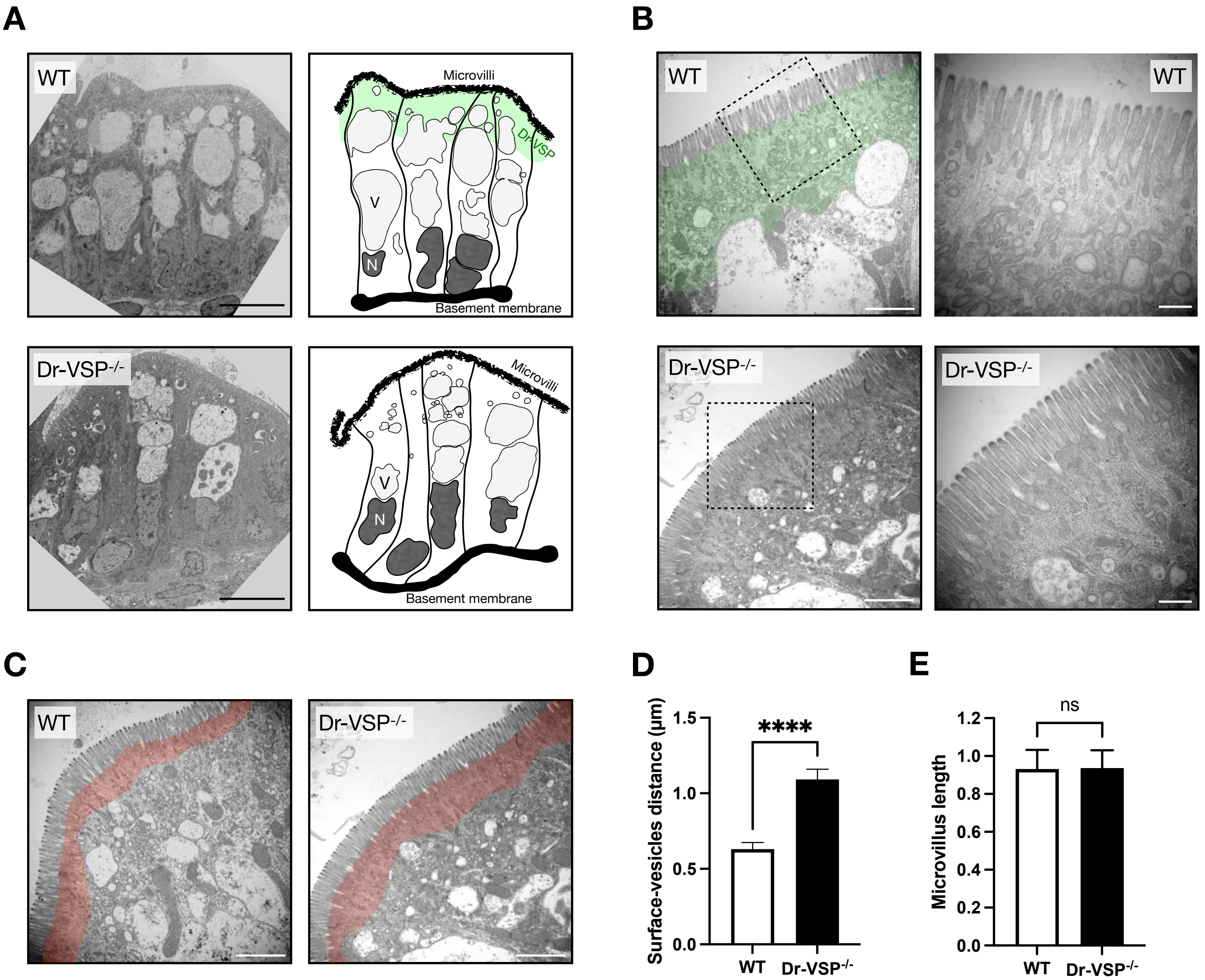

**Supplementary Figure S5: Morphological structures of zebrafish LREs under transmission electron microscopy.**

(A) Representative TEM images (left) of LREs from 14-dpf wild-type (top) and Dr-VSP$^{-/-}$ (bottom) zebrafish larvae; and their corresponding schematic illustration (right). Green color represents the subapical region upper to supranuclear vacuoles, where the positive immunofluorescence signal of Dr-VSP was observed in Figure 3, A and B. N, nucleus. V, vacuole. Scale bar = 10 µm.

(B) Representative TEM images of LREs from 14-dpf wild-type (top) and Dr-VSP$^{-/-}$ (bottom) zebrafish larvae, showing the ultrastructures of microvilli and subapical region. Key features of absorptive enterocytes are presented, including membrane invaginations at inter-microvillous spaces, cytoplasmic tubules and tubule-vacuole complexes, and numerous endocytic vesicles. In wild-type LRE, green color represents the subapical region upper to supranuclear vacuole, where immunofluorescence signal of Dr-VSP was observed in Figure 3, A and B. Scale bar = 2 µm (left) and 500 nm (right).

(C) Representative TEM images of LREs from 14-dpf wild-type (left) and Dr-VSP$^{-/-}$ (right) zebrafish larvae, showing the ultrastructures of microvilli and subapical region. Red color represents the area beneath apical surface that contains cytoplasmic tubules but few well-defined endocytic vesicles. The vesicles are more densely distributed in the areas deeper than the red region and are frequently associated with larger endosomal vacuoles.

(D) Inter-edge distance between apical surface and vesicle-dense area in wild-type and Dr-VSP$^{-/-}$ LREs, corresponding to the width of red shading areas shown in (C). Enterocytes, > 50 cells from 25 TEM images (magnification = 15000x) for each zebrafish line. Data were collected from 4 wild-types and 3 Dr-VSP$^{-/-}$ larvae. Average distances in each TEM image were analyzed using ImageJ macro.

(E) Microvillous length of wild-type and Dr-VSP$^{-/}$ LREs measuring at TEM level from the same samples as in (D). Error bars, means ± s.d.; ****P < 0.0001; unpaired Student's t-test; ns, no statistically significant difference.

#### Supplementary Figure S6

**A**

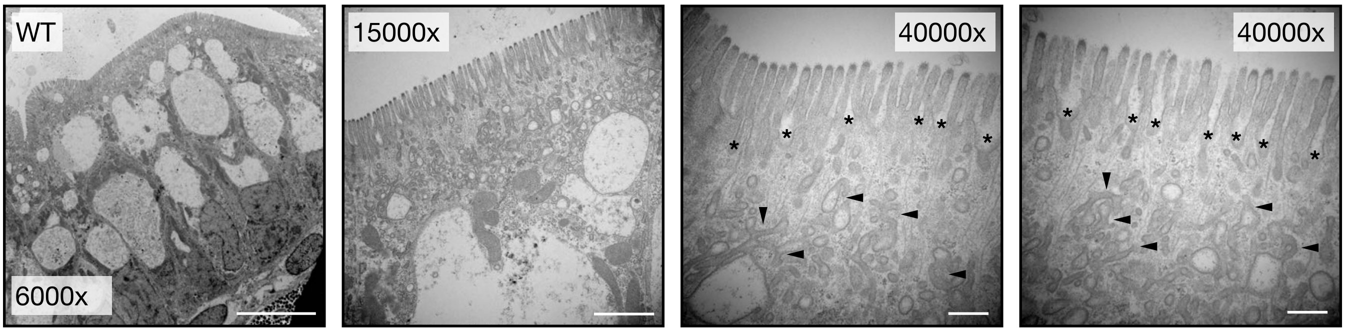


**B**

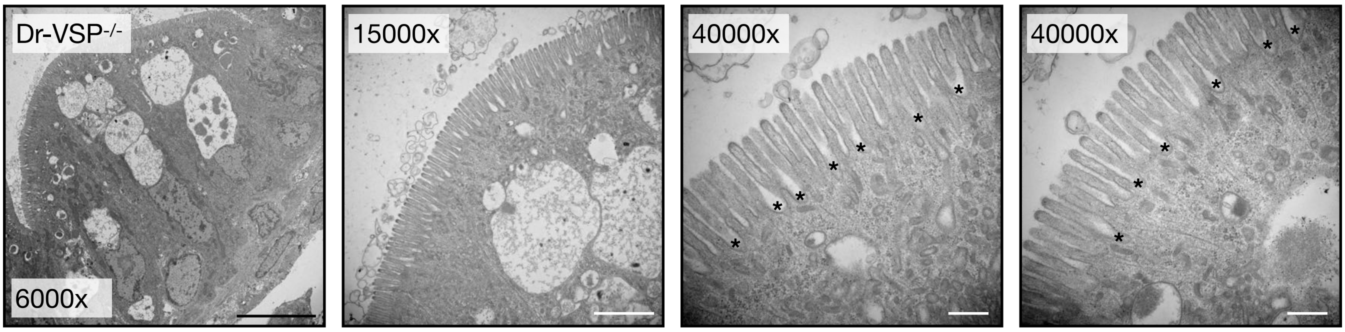

**Supplementary Figure S6: TEM image series of wild-type and Dr-VSP$^{-/-}$ LREs.**

Representative TEM image series of LREs from 14-dpf wild-type (top) and Dr-VSP$^{-/-}$ (bottom) zebrafish larvae. Large vacuoles are distributed in the supranuclear region but are fewer in Dr-VSP$^{-/-}$ LREs. Membrane invaginations at inter-microvillous spaces (*) can be observed at higher magnification. Branching cytoplasmic tubules and tubule-vacuole complexes (arrowheads) are more common in wild-type LREs. Scale bar at 6000x = 10 μm; at 15000x = 2 μm; at 40000x = 500 nm.

### Supplementary Figure S7

```
Hs-VSP          --------------------------------------------------------------------------------
Mm_VSP    1     MYGEKKSHLYLWMEHYGYDMPANIYKMYSQPSRKTDDANKKVSVSASRTIKLNGSTGYDTNEQITLITNGSSLSYPDEIK   80
Dr-VSP          --------------------------------------------------------------------------------
Cc-VSP          --------------------------------------------------------------------------------
On-VSP          --------------------------------------------------------------------------------
Ol-VSP          --------------------------------------------------------------------------------
Ci-VSP    1     M--------------EGFD--------------------------GSDFSPPADLVGVDG---------AVMRNVVD---   28
Ap-VSP    1     MC-------------QGIC-----------------------EIMADDTSPHRYLIDAEEENEPTLPLPTPLQDVPS---   41

Hs-VSP    1     ----------------------------------------------------MNESPQTNEFK-GTTEEAPAKESPHT--   25
Mm_VSP    81    SASYADPISTKAYTNDSSVYDPGGASSSTTLYELNSLSEVSKEIITQGESALLRDKEATSELKIPSTLQTQTSMSTNTLS   160
Dr-VSP    1     ------------------------------------------------------------------------------MT   2
Cc-VSP    1     ------------------------------------------------------------------------------MT   2
On-VSP    1     ------------------------------------------------------------------------------MS   2
Ol-VSP          ------------------------------------------------------------------------------M-
Ci-VSP    29    -----------VTINGDVTAPPKAAPRK-----SESVKKVHWNDVDQG----PSEKPETRQEE---RI------------   73
Ap-VSP    42    -----------TNVLAHSIFPPPVEPESKSDTSSGECKAVHL-VMTNGFLS------ANAEIGLDDTVPLGGRL------   97

                                                                       S1
Hs-VSP    26    -SEFKGAALVSPISKSMLE-RLSKFEVED--AEN--VASYDSKIKKIVHSIVSSFAFGIFGVFLVLLDVTLLLADLIFTD   99
Mm_VSP    161   LSDLSSDYQEEQMNKCKLN-QMSKLYDDD--ERTDIQKSYWNVVKKFVRILVSSVAFRIFGIFLVILDVFLVVVDLNVSE   237
Dr-VSP    3     SVHFNPGLDSKEVNGNSVK-EEAEVQIDDGKEETKDPDTMYHQVRKKITPFVMSFGFRVFGLVLIILDIIMVIVDLSLSE   81
Cc-VSP    3     SVHFNPGLDSKEVNGNGLK-EEAEVQIDDGREDTEDPDTMYHRVRKTIAPFVMSFGFRVFGLVLIIVDIILVIVDFSLSN   81
On-VSP    3     SVYFNPGSDSS-VNGNITKMENAKVEIDDGKEESALPDTLYYNLRKKIAPFVMSFGFRLFGVILIIVDFVLVIVDLSLPA   81
Ol-VSP    2     SVHFQPGSDAG-VNGNVAKMEDAKVEIDNGKEESALPDTLYHNIQRAITPFVMSFGFRLFGVVLIIVDFVLVIVDFSLAS   80
Ci-VSP    74    --------DIPEISGLWWGEN--EHGVDDGRME--IPTTGVGRVQFRVRAVIDHLGMRVFGVFLIFLDIILMIIDLSLPG   141
Ap-VSP    98    -LHDDPGENDEE---HWTENGHVQFELGDRAHVSVIRYPRLKKARMRTKRVVEHLAVRLIGVGLVIFDIIIVILDIVFNP   173

                            S2                                S3                          S4 →
Hs-VSP    100   SKLYIPLEYRSISLAIGLFFLMDVLLRVFVEGRQQYFSDLFNILDTAIIVIPLLVDVIYIFFDI----KLLRNIPRWTHL   175
Mm_VSP    238   KKIYIPLDYRSISLAIALFFLVDVLLRVSVEGRRRYFSDVLNTLDAVVIGVTVVVAVIYALYDK----HFLRDIPRLAVL   313
Dr-VSP    82    KSRDVGGAPETVSLVISFFFLIDVLLRVYVEGFKVYFSSKLNIVDACIVVITLVVTMIYAFSDF----SGASLIPRVVTF   157
Cc-VSP    82    GSHDVRRAMESVSLVISFFFLIDVLLRVYVEGFKVYFSSKLNIMDACIVVVTLVVTMIYAFTDL----SGASLIPRVVTF   157
On-VSP    82    KSRDAGNALEAVSLTISFFFLADVLLRVYVEGFKVYFSSKLNIIDACVVIVTLAVTMVYTFSDL----SGTSLIPRVVTF   157
Ol-VSP    81    KSREVAKALEAVSLSISFFFLADVLLRVYVEGFKVYFRSKLNIVDACVVVLTLVVTMVYTFSDL----SGVSLIPRVVNF   156
Ci-VSP    142   KSESSQSFYDGMALALSCYFMLDLGLRIFAYGPKNFFTNPWEVADGLIIVVTFVVTIFYTVLDEYVQETGADGLGRLVVL   221
Ap-VSP    174   QNIATMEAYDAISVAVWCYFLIEICLRIFAKG-KKFFRTKLDLLDLLIVTVTGSVTIVYVSVDL------TGSYLKLVVV   246

                →S4                 VSD-PD linker
Hs-VSP    176   VRLLRLIILIRIFHLLHQKRQLEKLMRRLVSENKRRYTRDGFDLDLTYVTERIIAMSFPSSGRQSFYRNPIEEVVRFLDK   255
Mm_VSP    314   LRPLRLLILIRILQLAHQKRQLERLTRKLVSGNKRRYKKDGFDLDLTYVTERIIAMSFPSSGRESFYRNPIKEVVRFLDT   393
Dr-VSP    158   LRSLRILILVRIFRLASQKRELEKVTRRMVSENKRRYQKDGFDLDLTYVTERVIAMSFPSSGKQALYRNPIREVVRFLDT   237
Cc-VSP    158   LRSLRILILVRVFRLASQKKELEKVTRRMVSENKRRYQKDGFDLDLTYVTDRVIAMSFPSSGKQALYRNPIREVARFLDT   237
On-VSP    158   LRFLRIIILVRVFRLAAQKKELEKVTRRMISENKRRYQKDGFDLDLTYVTDRVIAMSFPSSGKQAFYRNPIREVARFLDT   237
Ol-VSP    157   FRFLRIIILVRVFRLAAQKKELEKVTRRMISENKRRYQKDGFDLDLTYVTDRVIAMSFPSSGKQAFYRNPIKEVARFLDT   236
Ci-VSP    222   ARLLRVVRLARIF---YSHQQMKASSRRTISQNKRRYRKDGFDLDLTYVTDHVIAMSFPSSGRQSLFRNPIGEVSRFFKT   298
Ap-VSP    247   FRVLRIFLLVRLL---SERKHVSKATRKMVSQNKRRYREDDFDLDLTYITDRVIASSFPSSGSQAIYRNPIEEVARFLDT   323

                                                 WPD-loop                          Cx5R(T/S)
Hs-VSP    256   KHRNHYRVYNLCSERAYDPKHFHNRVSRIMIDDHNVPTLHEMVVFTKEVNEWMAQDLENIVAIHCKGGKGRTGTMVCALL   335
Mm_VSP    394   KHPNHYQVYNLCSERAYDPKHFHYRVRRIMIDDHNVPTLEEMLLFSKEVNNWMAQDPENVVAIHCKGGKGRTGTMVCACL   473
Dr-VSP    238   KHMDHYKVFNLCSEKGYDPKFFHYRVERVMIDDHNVPSLDDMLRYTACVRDWMAADSRNVIAIHCKGGKGRTGTMVCTWL   317
Cc-VSP    238   KHLDHYKVFNLCSEKGYDPKFFHYRVERVMIDDHNVPSLEDMLRYTACVRDWMAADSSNVIAIHCKGGKGRTGTMVCTWL   317
On-VSP    238   KHEGHYKVYNLCSEKGYDPQFFHYRVERVFIDDHNVPSLEDMLKYTASVREWMSADPQNIIAIHCKGGKGRTGTMVCTWL   317
Ol-VSP    237   KHEGHYKVFNLCSEKGYDPQHFHYRVERVFIDDHNVPALEDMLKFTASVREWMSADPQNVIAIHCKGGKGRTGTMVCTWL   316
Ci-VSP    299   KHPDKFRIYNLCSERGYDETKFDNHVYRVMIDDHNVPTLVDLLKFIDDAKVWMTSDPDHVIAIHCKGGKGRTGTLVSSWL   378
Ap-VSP    324   KHPDHYKVYNLCIERHYDESFFHNRVGRVLIYDHNVPRLKDLVAFCADARQWMEADNKNILFIHCKGGKGRTGLAVCAWL   403

                                       Gating loop
Hs-VSP    336   IASEIFLTAEESLYYFGERRTNKTHSNKFQGVETPSQNRYVGYFAQVKHLYNWNLPPRRILFIKRFIIYSI----RGDVC   411
Mm_VSP    474   IASEIVLNAKESLYFFGERRTDKSNSSKFQGIETPSQNRYVKYFEKLKINYQLTLPPKKVLVIKRLVVYSIHGVGKGDGS   553
Dr-VSP    318   IDSDQFESAQESLDYFGERRTDKSMSSKFQGVETPSQSRYVGYYEIMKNQYNRQLPPRKSLKIKSIRIHSIAGVGKGNGS   397
Cc-VSP    318   IDSDQFESAQESLDYFGERRTDKSMSSKFQGVETPSQSRYVGYYEIMKNQYNRQLPPQKSLKIKSIRIHSIAGVGKGNGS   397
On-VSP    318   IDSDQFESAQDSLEYFGERRTDKSQSSKFQGVETPSQSRYVGYYEIMKNKLNRQLPPPKSLRIKSIRIHSIAGVGKGNGS   397
Ol-VSP    317   IDSDQFESAQDSLEYFGERRTDKTRSSKFQGVETPSQSRYVGYYEVMKNKFNRQLPPPQSLRIKSLRIHSIAGVGRGDGS   396
Ci-VSP    379   LEDGKFDTAKEALEYFGSRRTDFEVGDVFQGVETASQIRYVGYFEKIKKNYGGQLPPMKKLKVTGVTITAIQGVGRGNGS   458
Ap-VSP    404   LESGVFTDTKECMGYFGVRRTDYTEGDMYQGVDTPSQSRYVDYFQKIKYSLGGHVPSPTPLTINTVRISGIQGVGKGDGR   483

                                                                            515loop (CBR3loop
Hs-VSP    412   DLKVQVVMEKKVVFSS--TSLGNCSILHDIETDKILINVYDGPPLYDDVKVQFFSS--NLPKYYDNCPFFFWFNTSFIQN   487
Mm_VSP    554   DLEVQIIMWQETVFSF--CNSRNCMIFHDPETDRAIINVFHCPALYDDVKVKFLSP--NLPKYYDDCPFFFWFHTSFIKN   629
Dr-VSP    398   DLKLKIIVKHELVFQCVCAKQHNCTVFPDTGSNAVVISLQDGPIVTGDVKVMFESSA-GLPKGYEDCPFYFWFNTSFVEN   476
Cc-VSP    398   DLKVKIIVKRELVFQCVCAKQQNCAVFPDTGNNAVVISLQEGPVVTGDVKVMFESSA-GLPKGYEDCPFYFWFNTSFVEN   476
On-VSP    398   DFKVKIVVRKELVFECVCAKQENCTVFPDVGNNAAVISLQNGPVVEGDVKVMFESSA-GLPKGYEDVPFYFWFNTSFIED   476
Ol-VSP    397   DLKVKIIVRGELVFQCVCAQQENCTVFPDAGSNAAVISLQNGPVVEGDVKVMFESSA-GLPKGYEDVPFYFWFNTSFVEH   475
Ci-VSP    459   DLSMQIVSERQEVLLCKFAEGYNCALQYDATDDCVTCEVKNCPVLAGDIKVRFMSTSKSLPRGYDNCPFYFWFNTSLVEG   538
Ap-VSP    484   DLWMEVHSDEVSLFSCDFRCNINCSVVYDFDMDTVIIQVKNSPAFSGDVQIKFKSSNKKIPKAYGECAFFFWFYTGFISD   563

Hs-VSP    488   NRLCLPRNELDNPHKQKAWKIYPPEFAVEILFGEK-----   522
Mm_VSP    630   NRLYLPRNELDNTHKPKTWKIYGEKFAVEVDFGEN-----   664
Dr-VSP    477   YRLFLSREELDNPHKPKTWDIYKEDFGVTLSFTEP-----   511
Cc-VSP    477   NSLYLSREELDNPHKSKTWDIYKEDFGVTLYFTDP-----   511
On-VSP    477   NKLFLPREELDNPHKPKTWDLYKEDFGVTMYFSESE----   512
Ol-VSP    476   NKLFFPREELDNPHKPKTWNLYKEDFGITVYFLPV-----   510
Ci-VSP    539   DHVTLKREEIDNPHKKKTWKIYRDNFTVKLTFSDAEDI--   576
Ap-VSP    564   NRLYLLRKDLDNPHKAEAWKVYRDTFAVDVTFGPVAESET   603
```

**Supplementary Figure S7:**

Multiple amino acid alignment of full-length VSP orthologs from human (*Homo sapiens*, Hs-VSP), mouse (*Mus musculus*, Mm-VSP), zebrafish (*Danio rerio*, Dr-VSP), common carp (*Cyprinus carpio*, Cc-VSP), Nile tilapia (*Oreochromis niloticus*, On-VSP), Japanese medaka (*Oryzias latipes*, Ol-VSP), sea squirt (*Ciona intestinalis* Type A, Ci-VSP), and Crown-of-thorns starfish (*Acanthaster planci*, Ap-VSP). Boxes indicate the putative transmembrane helices (S1 - S4), VSD-PD linker, WPD loop, CX5R(T/S), gating loop, 515 loop/CBR3 loop (Okamura et al., 2018). Amino acid sequences were retrieved from National Center for Biotechnology Information (NCBI) public database. Analyses were performed using online Constraint-based Multiple Alignment Tool (COBALT) (https://www.ncbi.nlm.nih.gov/tools/cobalt/)

### Supplementary Figure S8

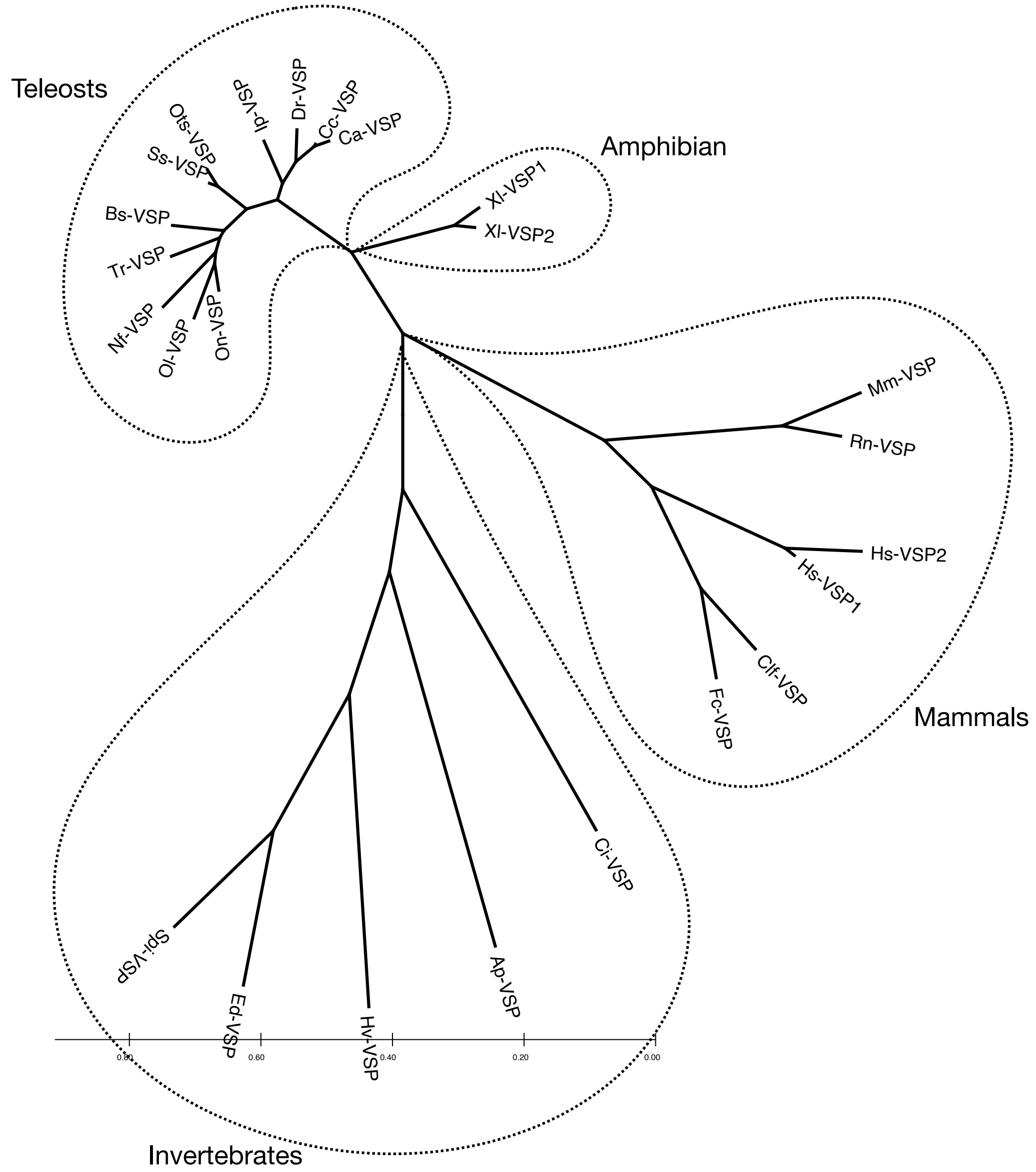

**Supplementary Figure S8:**

Molecular phylogenetic tree showing the phylogenic relationships among VSP orthologs from **mammals:** human (*Homo sapiens*, Hs-VSP1 and Hs-VSP2), mouse (*Mus musculus*, Mm-VSP), rat (*Rattus norvegicus*, Rn-VSP), dog (*Canis lupus familiaris*, Clf-VSP), cat (*Felis catus*, Fc-VSP); **amphibian:** American clawed frog (*Xenopus laevis*, Xl-VSP1 and Xl-VSP2); **teleosts:** goldfish (*Carassius auratus*, Ca-VSP), common carp (*Cyprinus carpio*, Cc-VSP), zebrafish (*Danio rerio*, Dr-VSP), channel catfish (*Ictalurus punctatus*, Ip-VSP), Chinook salmon (*Oncorhynchus tshawytscha*, Ots-VSP), Atlantic salmon (*Salmo salar*, Ss-VSP), Siamese fighting fish (*Betta splendens*, Bs-VSP), Japanese puffer (*Takifugu rubripes*, Tr-VSP), turquoise killifish (*Nothobranchius furzeri*, Nf-VSP), Japanese medaka (*Oryzias latipes*, Ol-VSP), Nile tilapia (*Oreochromis niloticus*, On-VSP); **invertebrates:** sea squirt (*Ciona intestinalis* Type A, Ci-VSP), crown-of-thorns starfish (*Acanthaster planci*, Ap-VSP), fresh-water polyp (*Hydra vulgaris*, Hv-VSP), sea anemone (*Exaiptasia diaphana*, Ed-VSP), and smooth cauliflower coral (*Stylophora pistillata*, Spi-VSP). Data were retrieved from the same source as Supplementary Figure S7, and analyzed using Molecular Evolutionary Genetics Analysis (MEGA) X Software (Pennsylvania State University, Pennsylvania, US).

#### Supplementary Figure S9

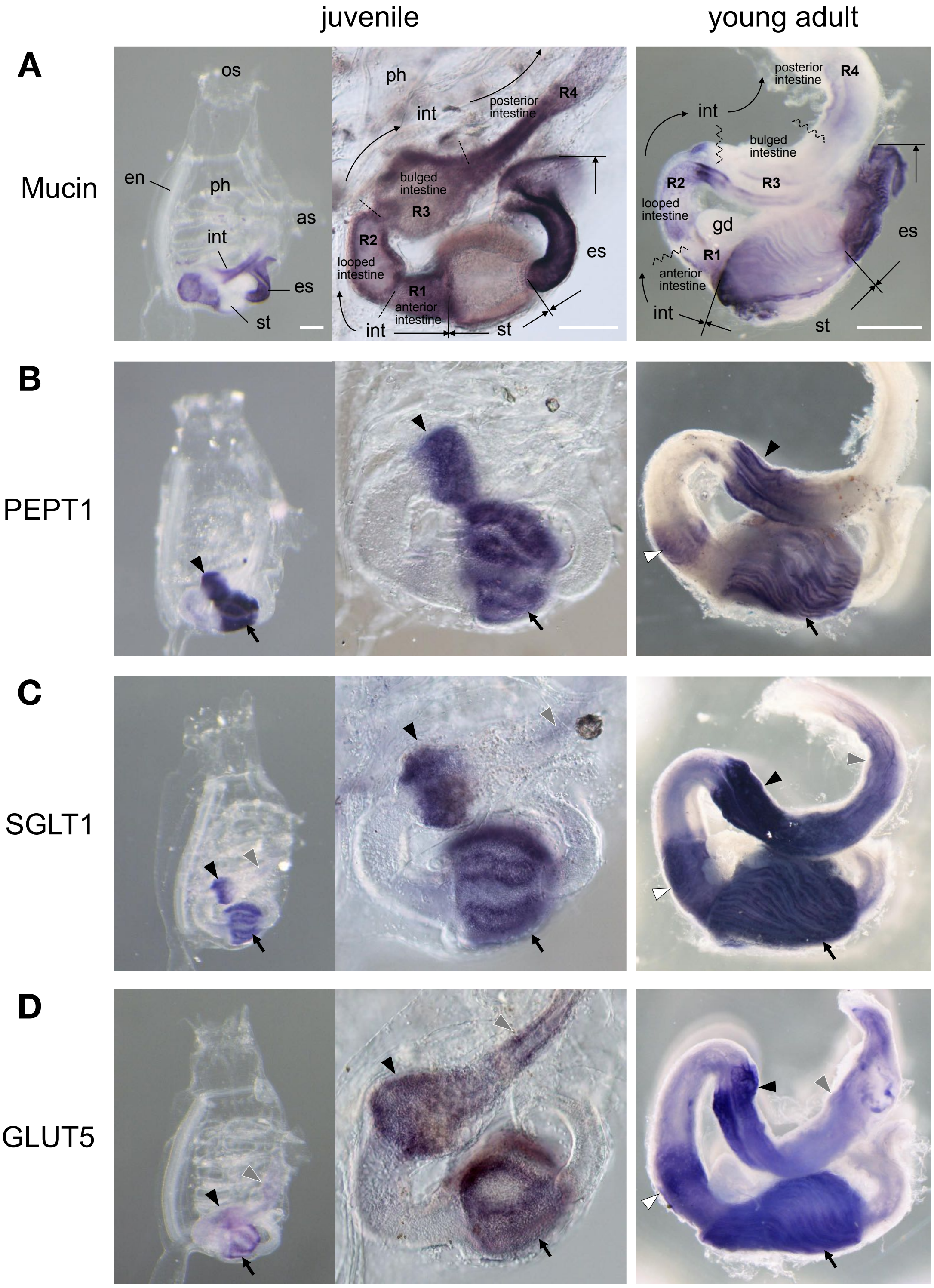

**Supplementary Figure S9: Morphological structure of ascidian intestine and expression profiles of intestine-related genes**

(A – D) Expression profiles of intestine-related genes in *Ciona* juvenile (1st column) and young adult (2nd column) revealed by WISH. In the juvenile, whole (left panel) and magnificent of the post-pharyngeal region of the digestive tract including esophagus, stomach, and intestine (right panel) were shown.

(A) Morphological structure of the digestive tract was noted on the expressions of Mucin gene. Digestive tract of *C. intestinalis* Type A was divided into oral siphon (os), pharynx (ph), esophagus (es), stomach (st), and intestine (int) (Chiba et al., 2004). The intestine was subdivided into four regions, R1 to R4 (Yoshida and Sasakura, 2012). Mucin gene was expressed broadly in the post-pharyngeal region of the digestive tract of the juvenile, but not expressed in the stomach and bulged-intestine (R3). In the young adult specimen, expression profile was similar to the juvenile, but expression signal was disappeared between anterior intestine (R1) and looped-intestine (R2) beside the developing gonad (gd).

(B) Absorption-related PEPT1 gene was invested as an example of the peptide transporter genes. Expression signals of the *PEPT1* were observed in stomach (arrow) and bulged-intestine (R3) of the mid-intestine (arrowhead) in both developmental stages of juvenile and young adult. Additional expression (white arrowhead) was appeared between anterior intestine (R1) and looped-intestine (R2) of the young adult intestine.

(C) Absorption-related SGLT1 gene was invested as a secondary active glucose transporter. Transcripts of the *SGLT1* were detected in stomach (arrow), bulged-intestine (R3) (arrowhead), and posterior-intestine (R4) (gray arrowhead) in juvenile and young adult specimens.

(D) GLUT5 gene was also assess as a facilitated transporter of the monosaccharide. Transcripts of the *GLUT5* were detected in stomach (arrow), bulged-intestine (R3) (arrowhead), and posterior-intestine (R4) (gray arrowhead) in juvenile and young adult specimens. Additional expression (white arrowhead) between anterior intestine (R1) and looped-intestine (R2) was also found in the young adult. Scale bars = 200 μm in (left) and 1 mm in (right). Additional abbreviations: as, atrial siphon; en, endostyle. Latest gene model IDs (HT version) of the *Ciona Mucin*, *PEPT1*, *SGLT1*, and *GLUT5* are KY.Chr2.2226, KY.Chr13.370, KY.Chr10.1303, and KY.Chr7.810, respectively (Satou et al., 2019)

### Supplementary Movie S1

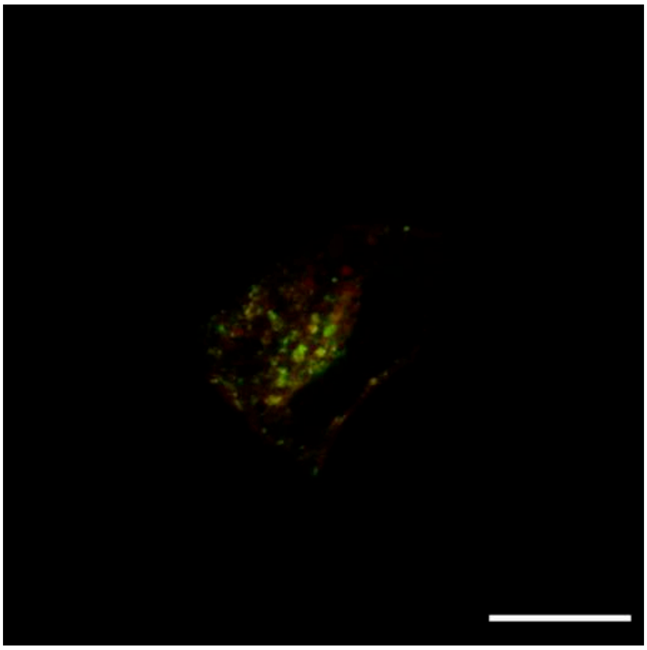

**Supplementary Movie S1: Dr-VSP is localized at the endosomal membranes of early endosomes.**

MDCKII cells co-expressing Dr-VSP-mCherry and Rab5-EGFP for early endosomes. Time-lapse images were acquired from the same samples as presented in Figure 4B. Red, Dr-VSP. Green, Rab5. Scale bar = 20 μm.

**Supplementary Movie S2**

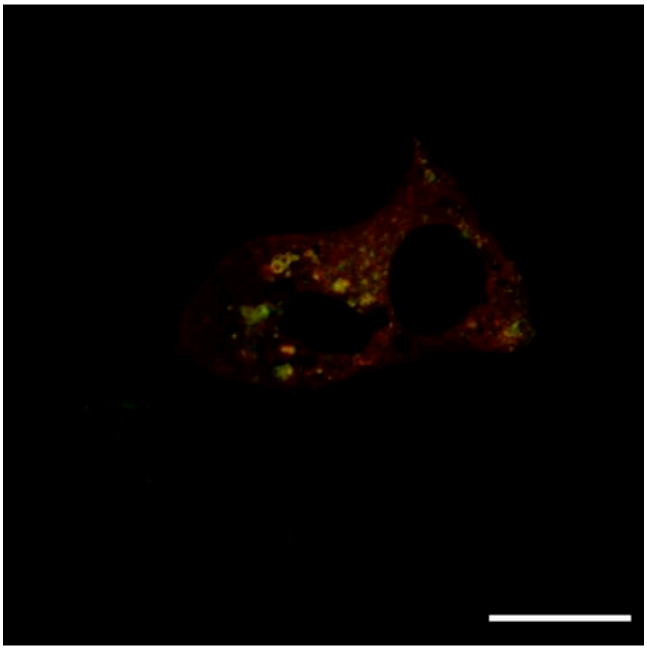

**Supplementary Movie S2: Dr-VSP is localized at the endosomal membranes of recycling endosomes.**

MDCKII cells co-expressing Dr-VSP-mCherry and Rab11-EGFP for recycling endosomes. Dr-VSP is localized at the endosomal membranes of recycling endosomes. Time-lapse images were acquired from the same samples as presented in Figure 4B. Red, Dr-VSP. Green, Rab11. Scale bar = 20 μm.
