## Supplementary material for "Voltage-Sensing Phosphatase (VSP) Regulates Endocytosis-Dependent Nutrient Absorption in Chordate Enterocytes": DrVSP Supplementary Table S1

**Supplementary Table S1: RT-PCR Primers used in this study**

| Target gene | Primer (5' – 3') | |
|---|---|---|
| *vsp* | Sense | CAGAGCAGGTATGTTGGCTA |
| | Antisense | CAGCACTGGACTCAAACATGA |
| *β-actin* | Sense | GGTATGGAATCTTGCGGTAT |
| | Antisense | GGTATGGAATCTTGCGGTAT |
