## Supplementary material for "Voltage-Sensing Phosphatase (VSP) Regulates Endocytosis-Dependent Nutrient Absorption in Chordate Enterocytes": DrVSP Supplementary Table S2

**Supplementary Table S2: CRISPR target sequences of *vsp* and PCR primers used in this study**

| Target sequence | | Primer (5' – 3') | |
|---|---|---|---|
| T1 | GGACATCATTATGGTCATTG**TGG** | Sense | GGTGAGTGGCATACTGGAATTT |
| | | Antisense | TCCACATAAACACGGAGCAATA |
| T2 | GAGAAGAGTCGTGATGTTGG**AGG** | Sense | TTATGTTTGCAGTGTTTTTGGC |
| | | Antisense | ACATCATGTGCTGAAATCAAGG |
| T3 | TGCCACTCACCGAAAACCAA**AGG** | Sense | TCTGATTTGTTTTGCAGTCAGG |
| | | Antisense | CCAACATCACGACTCTTCTCAG |

Bold letters at the end of each target sequence indicate the protospacer adjacent motif (PAM) sequence.
