## Supplementary material for "Voltage-Sensing Phosphatase (VSP) Regulates Endocytosis-Dependent Nutrient Absorption in Chordate Enterocytes": DrVSP Supplementary Table S3

**Supplementary Table S3: Key resources table**

| REAGENT or RESOURCE | SOURCE | IDENTIFIER |
|---|---|---|
| **Antibodies** | | |
| Mouse monoclonal Anti-Zebrafish VSP/TPTE (clone N432/21) | UC Davis/NIH NeuroMab Facility | Cat# 73-485; RRID: AB_2716253 |
| Rabbit monoclonal Anti-Sodium Potassium ATPase | Abcam | Cat# ab76020; RRID: AB_1310695 |
| Acti-stain™ 488 Phalloidin | Cytoskeleton, Inc. | Cat# PHDG1 |
| Alexa Fluor™ 594 Phalloidin | Invitrogen | Cat# A12381; RRID: AB_2315633 |
| **Chemicals, peptides, and recombinant proteins** | | |
| Dextran, Alexa Fluor™ 488-tagged; 10,000MW | Molecular Probes, Inc. | Cat# D22910 |
| mCherry solution | This study | N/A |
| TRIzol™ LS reagent | Invitrogen | Cat# 10296010 |
| Anti-Digoxigenin-AP, Fab fragments | Roche | Cat# 11093274910 |
| BCIP/NBT solution | Wako | Cat# 022-16231 |
| BCIP/NBT solution | Roche | Cat# 11681451001 |
| Agarose, Ultra-low Gelling Temperature | Sigma-Aldrich | Cat# 9012-36-6 |
| **Critical commercial assays** | | |
| SuperScript™ III First-Strand Synthesis System | Invitrogen | Cat# 18080051 |
| PCR DIG Probe Synthesis Kit | Roche | Cat# 11636090910 |
| BigDye™ Terminator v3.1 Cycle Sequencing Kit | Thermo Fisher | Cat# 4337457 |
| Lipofectamine™ 3000 transfection reagent | Invitrogen | Cat# L3000015 |

| | | |
|---|---|---|
| **Experimental models: Cell lines** | | |
| MDCK-II Cell Line canine | ATCC | CRL-2936 |
| **Experimental models: Organisms/strains** | | |
| Zebrafish/ RIKEN wild-type | RIKEN Brain Science Institute | N/A |
| Zebrafish/ *Tg(vsp:EGFP)* | This study | N/A |
| Zebrafish/ Dr-VSP$^{-/-}$ | This study | N/A |
| *Ciona intestinalis* Type A | This study | N/A |
| **Oligonucleotides** | | |
| RT-PCR Primers used in this study | This study; Table S1 | N/A |
| CRISPR target sequences of *vsp* and PCR primers used in this study | This study; Table S2 | N/A |
| **Recombinant DNA** | | |
| Mbait-hsp70 promoter-EGFP-polyA | Kimura et al.[29] | N/A |
| mCherry-Dr-VSP | This study | N/A |
| EGFP-Rab5 | This study | N/A |
| EGFP-Rab11 | This study | N/A |
| **Software and algorithms** | | |
| ImageJ | NIH Image | http://imagej.nih.gov/ij |
| GraphPad Prism 8 | GraphPad Software | https://www.graphpad.com |
| MEGAX | MEGA | https://www.megasoftware.net |

| | | |
|---|---|---|
| COBALT | NIH NCBI | https://www.ncbi.nlm.nih.gov/tools/cobalt |
| iDEP v0.92 | Ge et al. [54] | http://bioinformatics.sdstate.edu/idep92 |
